# A diverse community constitutes global coccolithophore calcium carbonate stocks

**DOI:** 10.1101/2025.09.11.675535

**Authors:** Joost de Vries, Fanny M. Monteiro, Alex J. Poulton, Nicola A. Wiseman, Levi J. Wolf

**Affiliations:** School of Geographical Sciences, University of Bristol, BS8 1SS, UK; Centre for ice, Cryosphere, Carbon and Climate (iC3), Department of Geosciences, UiT The Arctic University of Norway, 9019, Tromsø, Norway; The Lyell Centre for Earth and Marine Science, Heriot-Watt University, Edinburgh, EH14 4BA, UK

**Author notes:** **Correspondence:** Joost de Vries.

## Abstract

Coccolithophores are diverse calcifying plankton, yet most research has focused on a single species, *Gephyrocapsa huxleyi*, with the global contributions of other species hitherto unexplored. Since coccolithophores account for the majority of marine calcium carbonate (CaCO_3_) production, this narrow focus risks biasing our understanding of CaCO_3_ cycling, as other species differ in their distributions, CaCO_3_ production and response to climate change. Using a global, species-resolved machine-learning approach, we show that *G. huxleyi* contributes only about 7% of estimated coccolithophore CaCO_3_ stock, while a morphologically and functionally diverse assemblage dominates. Since stock contributions are a good proxy for contribution to production, our findings challenge the view that *G. huxleyi* underpins CaCO_3_ cycling and show that lab and in situ datasets centred on this species capture only a small fraction of coccolithophore calcification. Our work identifies key species and regions to guide future laboratory, in situ, and modelling efforts, laying the groundwork for more realistic representations of CaCO_3_ cycling under climate change.

## 1 Introduction

Coccolithophores are major contributors to the ocean carbon cycle, affecting atmospheric carbon dioxide and, consequently, the global climate (Balch, 2018; Neukermans et al., 2023; Monteiro et al., 2016). Despite being microscopic (Figure 1), coccolithophores grow in large quantities in our oceans. They contribute to global oceanic primary production via photosynthesis (up to 10 %) (Poulton et al., 2007) and dominate global oceanic calcium carbonate (CaCO_3_) production through the formation of coccoliths (Balch and Mitchell, 2023; Neukermans et al., 2023; Ziveri et al., 2025). These processes are critical for regulating global climatic changes over different timescales. On short timescales, coccolithophore activity affects air-sea CO_2_ fluxes by altering ocean alkalinity (Kwon et al., 2024; Krumhardt et al., 2020; Planchat et al., 2024), while on longer timescales, coccolithophores facilitate CaCO_3_ accumulation in marine sediments that help buffer large atmospheric CO_2_ variations (Ridgwell and Zeebe, 2005; Boudreau et al., 2018).

**Figure 1.**
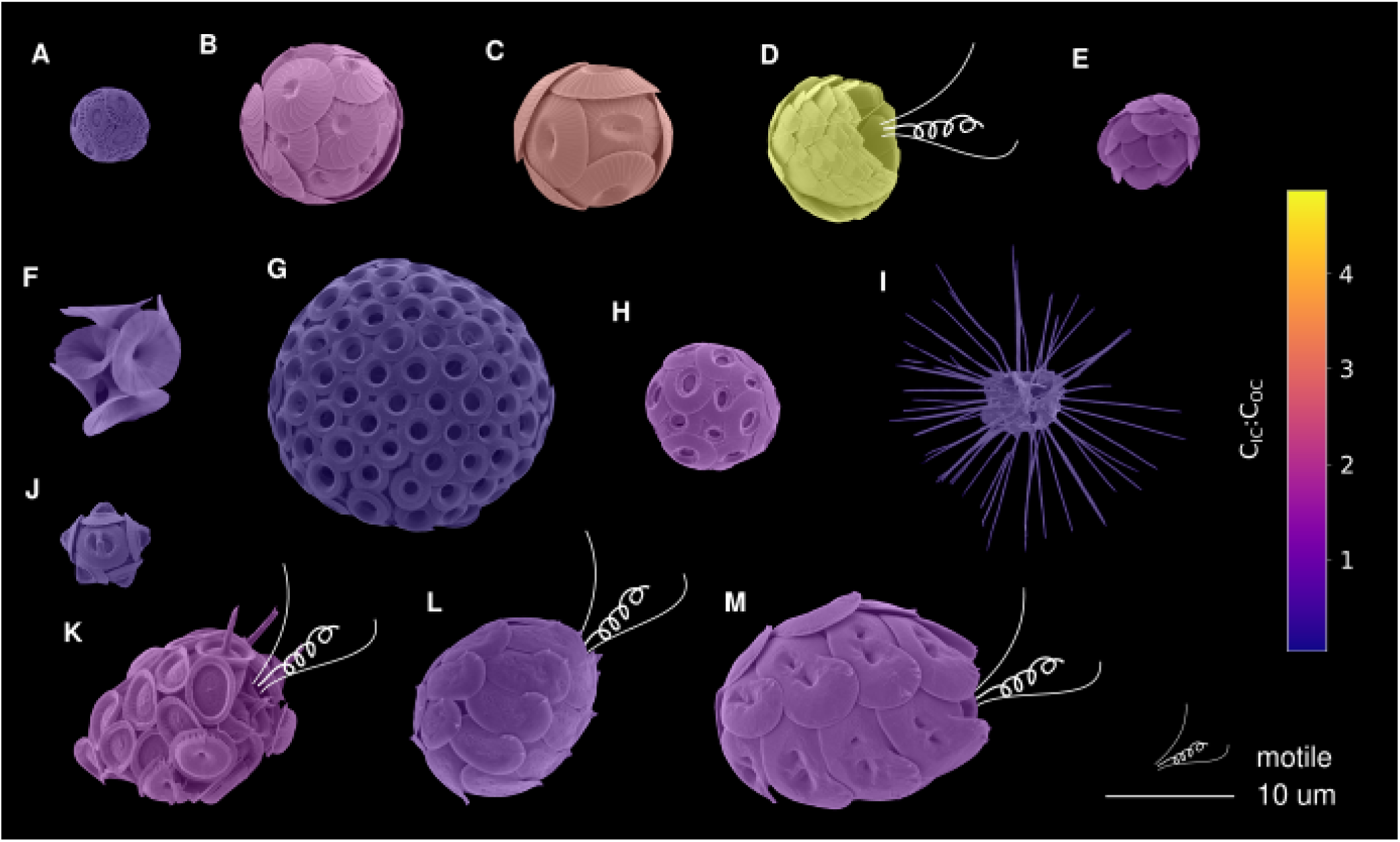
The morphology of notable coccolithophore species included in this study. **(A)** Gephyrocapsa huxleyi. **(B)** Calcidiscus leptoporus. **(C)** Coccolithus pelagicus. **(D)** Florisphaera profunda. **(E)** Oolithotus antillarum. **(F)** Umbellosphaera irregularis. **(G)** Umbilicosphaera sibogae. **(H)** Umbilicosphaera hulburtiana. **(I)** Palusphaera vandelii. **(J)** Gephyrocapsa oceanica. **(K)** Syracosphaera pulchra. **(L)** Helicosphaera hyalina. **(M)** Helicosphaera carteri. The Scanning Electron Microscopy (SEM) images are coloured by the ratio between the cellular organic and inorganic carbon (C_IC_:C_OC_) contents. C_IC_:C_OC_ data was retrieved from the CASCADE database (de Vries et al., 2024). For images of all 58 species included in this study see Figure A1. SEM images by Jeremy Young and acquired from Nannotax3 (Young et al., 2022).

About 250-300 coccolithophore (morpho)species live in the ocean (Young et al., 2022), displaying remarkable diversity in traits such as cell size, motility, morphology and calcite content (Figure 1 and Figure A1) (Monteiro et al., 2016). Since these traits influence distributions and their response to environmental changes (Litchman and Klausmeier, 2008), coccolithophore responses to climate change are expected to vary between species. Despite this, most research has focused on *Gephyrocapsa huxleyi* (formerly *Emiliania huxleyi*), one of the most abundant and widespread coccolithophore species (O’Brien et al., 2013; de Vries et al., 2021). For instance, global estimates of coccolithophore distribution, stocks and production primarily rely on satellite algorithms that are heavily calibrated to *G. huxleyi* reflectance (Balch and Mitchell, 2023; Hopkins et al., 2019). Similarly, climate models that include coccolithophores often treat them as a single functional group, parameterised primarily on *G. huxleyi* characteristics (Le Quéré et al., 2016; Seifert et al., 2022; Krumhardt et al., 2019, 2020).

Despite the high abundance of *G. huxleyi*, regional scale studies show other species dominate coccolithophore CaCO_3_ standing stocks, production and export fluxes (Daniels et al., 2016; Rigual Hernández et al., 2020). Larger or more heavily calcified species drive regional CaCO_3_ production and export fluxes, such as *Coccolithus pelagicus*, *Calcidiscus leptoporus* and *Florisphaera profunda* (Figure 1) (Daniels et al., 2016; Rigual Hernández et al., 2020; Guerreiro et al., 2017). A global study using sediment trap data also revealed that species other than *G. huxleyi* dominate deep-sea fluxes (Ziveri et al., 2007).

Although standing stock represents the instantaneous balance between production, loss, and removal and does not directly quantify carbonate production, it remains a useful proxy for comparing species contributions to pelagic CaCO_3_ production. Production will vary with growth and calcification rates (Krumhardt et al., 2017), but these differences do not follow simple assumptions about “fast” versus “slow" growing taxa. For instance, *G. huxleyi* (a presumed fast growing species) and *Coccolithus pelagicus* (a presumed slow growing species) have been observed to exhibit broadly comparable growth under identical temperature and light conditions (Daniels et al., 2014). As such, most studies converting coccolithophore stocks to contributions reliably use a single growth rate for all species (Ziveri et al., 2023; Han et al., 2025; Kruijt et al., 2026). Thus, while stock-based comparisons should not be interpreted as direct measures of production, they provide meaningful insight into species-specific functional importance.

Since high cellular calcification increases sensitivity to pH (Gafar et al., 2019; Kottmeier et al., 2022), heavily calcified species may be more sensitive to changes in carbonate chemistry than the more lightly calcified *G. huxleyi*. Furthermore, traits like motility and cell size will influence sensitivity to climate change impacts such as warming and stratification (Litchman and Klausmeier, 2008; Ziveri et al., 2025). Thus, a focus on a small, non-motile and lightly calcified species may misrepresent coccolithophores’ vulnerability to climate change.

While previous research suggests that species other than *G. huxleyi* might contribute significantly to coccolithophore stocks, no global study has characterized species-specific contributions. To address this, we employ machine learning algorithms to generate statistically robust global estimates of biomass and inorganic carbon stocks of the most frequently occurring coccolithophore species. This analysis uses the new water column coccolithophore abundance dataset CASCADE and associated species-specific estimates of cellular inorganic carbon content (de Vries et al., 2024; Sheward et al., 2024). This study presents the first global, species-specific assessment of coccolithophore carbon stocks, providing a far more accurate understanding of their role in the carbon cycle.

## 2 Materials and Methods

We built global spatial-temporal (Latitude, Longitude, Depth, Month) species distribution models (SDMs) of coccolithophores to determine the key species contributing to coccolithophore cellular organic carbon (C_OC_) and cellular inorganic carbon (C_IC_) standing stocks. To build our SDMs, we used a Python ensemble-based machine learning pipeline (A_BIL_._PY_, v2025.08.06 (de Vries et al., 2025b, 2021)), which was trained on coccolithophore species abundance data provided in the CASCADE dataset (de Vries et al., 2024). The resulting machine learning models were used to fill in gaps in abundance observations based on interpolated monthly 1-degree gridded environmental data. The predicted abundances were converted to stocks of organic carbon (C_OC_) and inorganic carbon (C_IC_) using species-specific conversion factors from the CASCADE data set (de Vries et al., 2024). Finally, we calculated confidence intervals for our estimates by leveraging the uncertainty quantification capabilities of ensemble-based machine learning methods. These uncertainty measures were then applied to the globally integrated stock estimates, as well as the species-specific contributions to this estimate.

### 2.1 Training and prediction data

#### 2.1.1 Coccolithophore observations

We used coccolithophore microscopy abundance data in the CASCADE dataset (de Vries et al., 2024), which collects measurements of the abundance of coccolithophore species across 6,166 water samples taken from 1964 to 2019 (Figure 2). We focused our analysis on species occurring in at least 5 % of the locations (*>* 100 observations, 58 species). We excluded rarer species, as such species are unlikely to be major contributors to global carbon standing stocks. Furthermore, we excluded the species *Oolithotus fragilis* from our analysis as this species is easily confused with *Coccolithus leptoporus* in light microscopy measurements (Young et al., 2022) thus potentially leading to double accounting.

**Figure 2.**
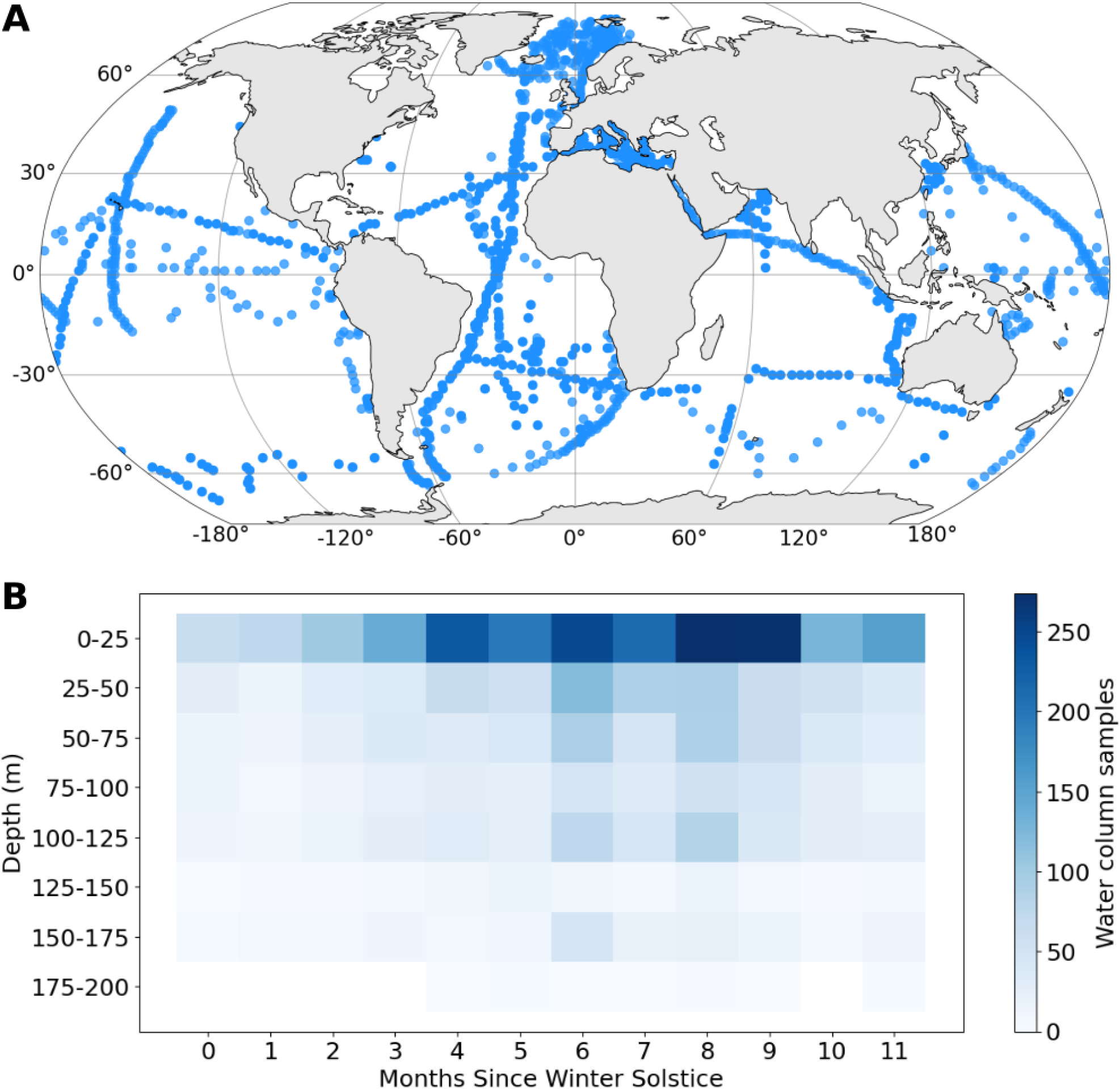
Sample distribution of the coccolithophore abundances from the CASCADE database used as training data in this study. **(A)** Latitude-longitude distributions of observations. **(B)** Depth-time distributions. Note that for the temporal plot the months since winter solstice was calculated to account for hemispherical differences in phenology.

For the modelled species we used all available oceanic species-specific abundance observations in CASCADE which were gridded to a monthly, global 1-degree grid, with 5 m depth levels up to 200 meters deep. We included data from both Scanning Electron Microscopy (SEM) and Light Microscopy (LM) images.

#### 2.1.2 Environmental data

To build predictive models of coccolithophore distribution, we use interpolated monthly mean grids of temperature, phosphate, nitrate, silicic acid, oxygen, photosynthetically active radiation (PAR), dissolved inorganic carbon (DIC) and total alkalinity (TA). These features were selected based on mechanistic processes to avoid spurious correlations. From the World Ocean Atlas 2023 (Reagan et al., 2024), we used temperature, nitrate, silicate, phosphate, and oxygen. PAR data was acquired from Castant et al., 2024 (Castant et al., 2024), and NNGv2 (Broullón et al., 2019, 2020) provided data for DIC and TA.

#### 2.1.3 Organic and inorganic carbon

We converted individual species abundance into organic carbon (C_OC_) and inorganic carbon (C_IC_) biomass (pg C cell l*^−^*^1^) using data provided in the CASCADE dataset (de Vries et al., 2024). The dataset includes species-specific C_OC_ and C_IC_ estimates, which can vary 25-fold between species (de Vries et al., 2024). However, estimates in the CASCADE dataset do not consider inter-specific environmental controls of cellular C_OC_ and C_IC_ content, which can vary depending on environmental variables such as temperature, irradiance, and carbonate chemistry (de Vries et al., 2024; Gafar et al., 2019).

### 2.2 Model configuration

#### 2.2.1 Model description

To predict the distribution of each species, we utilize a 2-phase regressor to predict species abundance in each 1 degree × 1 degree × 5 m x 1 month grid point (‘voxel’) based on environmental conditions within the voxel (Figure A2). In this model setup, there are two steps: 1) a classifier predicts if the species is present or absent in each voxel based on environmental data. 2) if the species is predicted to be present, a regressor then predicts the concentration of biomass. We used a 2-phase model as it allowed us to predict absences and biomass concentrations separately, which was necessary due to the lack of explicit absence data (see section below). The two-phase model structure allows us to introduce pseudo-absences, or artificial measurements that indicate voxels where coccolithophores are not found, to calibrate biomass models more effectively. This is detailed in the next section.

#### 2.2.2 Absence prediction

The coccolithophore abundance dataset used to train our models (CASCADE) contains no zero observations. This is because the absence of observations (“Not a Number”, NAN values) does not necessitate the species’ absence (0 values) for a given sample. However, since the algorithms included in our analysis do not extrapolate, without absences in the data, non-zero values cannot be predicted without including pseudo-absences. To generate pseudo-absences in the dataset, we randomly introduce zero values based on area-of-applicability (AOA) estimates, introducing absences only outside of the AOA for a given species. To estimate AOA, we follow a strategy similar to Meyer and Pebesma (2021) (Meyer and Pebesma, 2021). For each dataset, a cutoff distance was determined using the 90th percentile inter-fold feature distance among training instances. Then, test samples further than this cutoff were determined to be “inapplicable” samples for the model, since their environmental conditions are quite dissimilar from the other observed environmental conditions.

To determine the optimal pseudo-absence ratios (i.e. ratio of artificial absence measurement to real presence measurement), full-ensemble runs were conducted with five different ratios: 1:0, 1:1, 1:2, 1:3, 1:4 and 2:1 (presence:absence). For each ratio, we estimated the summary statistics R^2^, relative Root Mean Squared Error (rRMSE), and relative Mean Average Error (rMAE) and total and species specific monthly integrated C_IC_ and C_OC_ (Figure A3). As pseudo-absence ratios increase, carbon tends to decrease. Thus, plots of pseudo-absence prevalence vs carbon totals were used as “scree” or “elbow” plots converging to an asymptote (Figure A3) to determine the minimal level of pseudo-absences required for effective estimation. For the ideal pseudo-absence ratio, the minimum number of pseudo-absences within the uncertainty bounds of the asymptote was selected, as AOA might introduce pseudo-absence in low biomass regions due to sampling biases. Based on this, a pseudo absence ratio of 1:1 was chosen.

#### 2.2.3 Voting ensembles

To merge estimates from the different types of algorithms, voting ensembles weighted based on their Mean Average Error (MAE) were used. Voting ensembles are a method in which the predictions from multiple classifier or regression models are combined to make a final prediction. This improves predictive performance as aggregating predictions from various sources can lead to better results (Géron, 2022). For our voting ensembles, three machine-learning algorithms were included: bagged K-Nearest Neighbors (b-KNN), Random Forests (RF), and Extreme Gradient Boosting (XGB). Each of these models is an ensemble of weak learners, which only see a subset of the data and the features, which prevents over-fitting when combined with cross-validation. All models were configured using _SCIKIT_-_LEARN_ (Varoquaux et al., 2015), except for gradient boosting, for which the XGBoosting algorithm package (Chen and Guestrin, 2016) was also used. A brief conceptual description of each algorithm is provided below.

##### Random Forest

The Random forests algorithm combines multiple decision trees; each trained on a distinct subset of the data set (Breiman, 2001). Furthermore, each decision tree uses a random subset of features (environmental data), mitigating bias towards specific features (Breiman, 2001). The final prediction is made by aggregating the predictions of individual trees.

##### XGBoost

Like random forests, the XGBoost (Extreme Gradient Boosting) algorithm combines multiple decision trees, each trained on a subset of the data (Chen and Guestrin, 2016). However, XGBoost works by iterative training and adding decision trees to the model, correcting for the errors made by the previous trees by increasing the weights of poorly predicted samples (Chen and Guestrin, 2016).

##### KNN

Finally, the K-Nearest Neighbors (KNN) algorithm calculates the distance between an input sample and its neighbours in the feature space (here, the environmental data) (Cover and Hart, 1967). Here, we extended the KNN algorithm with an ensemble bagging step (b-KNN), randomly splitting the data before training multiple KNN algorithms. A final prediction is then made by aggregating the predictions of individual KNN models

#### 2.2.4 Hyperparameter optimization

To optimize model performance, we conducted hyperparameter tuning for each model configuration using a ten-cross-fold validation (CV). This optimization was based on balanced accuracy for the classifiers and R^2^ values for the regressors. Hyperparameter tuning is a process that tries different model configurations (e.g., the maximum number of samples provided for each tree in the random forest). Then, it selects the best parameters based on model performance. During CV, a data ‘fold’ is generated by randomly splitting the data into test (10 %) and training data (90 %). This is done ten times to create ten random folds. For each fold, the model is trained on the training data and then validated on the test data. Mean out-of-sample ("test") error metrics can then be calculated by taking the average of the folds, which is then used to pick the best model configuration. The hyperparameters explored are provided and described in Table A1.

#### 2.2.5 Transformations

For all models, predictive performance was compared with and without log transformation of the target variable (species abundance); the best model for each species was then picked based on R^2^. Log transformation reduces the relative importance of high-value abundance observations. For species with outlier values, log transformation can thus improve model performance by being more representative of the bulk of the data but performs worse if high values are not outliers. To ensure accurate estimation of model performance when log transformations were applied, transformations were back-transformed before validation using _SCIKIT_-_LEARN_ (Varoquaux et al., 2015).

#### 2.2.6 Presence threshold

To determine the presence probability threshold at which the species was classified as present, we used Receiver Operating Characteristic (ROC) curve analysis. This method evaluates the trade-off between true positive rates (sensitivity) and false positive rates (1-specificity) across various probability thresholds. The optimal threshold was identified using the Youden Index, which maximizes the difference between sensitivity and specificity, providing a balanced and statistically robust criterion for species presence.

### 2.3 Uncertainty quantification

#### 2.3.1 Confidence intervals

Ensemble-based machine learning methods such as the RF, b-KNN used in this study, offer inherent uncertainty quantification. While XGB can be adapted to provide similar estimates by adding a bagging step. In these methods, each ensemble member generates a prediction, and these predictions can be utilized to estimate 95th percentile confidence intervals based on the prediction distribution.

Here, we separate the uncertainty quantification for each phase of the model, by considering the distributions of the classifier and regression models separately. However, to make our final predictions, the two models are combined using values from the summary statistic, and the previously determined presence threshold.

#### 2.3.2 Model extrapolation

To identify regions of high uncertainty due to limited sampling coverage we use Area of applicability (AOA) estimates to define the voxels (Latitude x Longitude x Depth x Time) which fall outside of our training data. To visualize this data along reduced dimensions (i.e. Latitude x Longitude, Latitude x Depth, and Time) voxel biomass values found applicable (applicable total) are estimated for each species and then summed. These values are then divided by total biomass values of all species (full total). Finally, the proportion of biomass for a given location is multiplied by 100 to derive the percentage of biomass that is extrapolated. For the extrapolation estimate calculated along time only, calendar month was converted to the number of months since the winter solstice to account for hemispherical differences in the seasonal cycles.

Like previously, we used the method described in Meyer and Pebesma (2021) (Meyer and Pebesma, 2021) for AOA estimates, however instead of using percentiles, we follow the author’s original recommendation and used the Tukey rule, setting the cutoff at 1.5 times the inter-quartile range (IQR) above the upper hinge (75th percentile) for the train data dissimilarity index.

### 2.4 Post processing

#### 2.4.1 Biome contributions

To aid interpretation and to estimate relative importances of key oceanographic regions we used core biomes defined in Fay and McKinley 2014 (Fay and McKinley, 2014). These biomes are defined based on sea surface temperature, spring/summer chlorophyll-a concentration, ice fraction, and maximum mixed layer depth (Fay and McKinley, 2014). For conciseness we merged the 17 biomes defined based on ocean basin or hemisphere into 5 superbiomes (‘ice’, ‘subpolar’, ‘subtropical seasonally stratified’, ‘subtropical permanently stratified’ and ‘equatorial’).

#### 2.4.2 Photic zone contributions

Photic zones were defined based on ecological specification specified in Poulton et al., 2017 (Poulton et al., 2017). This classification splits the water column into three different zones and includes the Upper Euphotic Zone (*>* 10 % surface irradiance), the Lower Euphotic Zone (1-10 %) and the Sub-Euphotic Zone (*<* 1 %). Percentage irradiance values were estimated based on PAR data from Castant et al., 2024 (Castant et al., 2024).

#### 2.4.3 Percentage contributions

To estimate percentage contributions, for a given biome, photic zone or species, the confidence intervals derived from the ensembles were used for sample reconstruction. These reconstructed distributions were then randomly sampled (n=10,000), and the resulting contribution to total stock was then estimated for each sample. The original biomass values were assumed to be Gamma distributed and shape and rate parameters were inferred on the 95th percentile quantiles based on the moments of these Gamma-distributed variates, after translating their parameters from scale-rate to location-scale expressions.

#### 2.4.4 Cumulative contributions

To estimate the number of species that are needed to account for 80 % of carbon stocks, we ranked the species based on their integrated contribution to the stock (C_OC_ or C_IC_). Cumulative contributions were then used to estimate the number of species required to reach the 80 % threshold.

#### 2.4.5 Diversity indices

Diversity was estimated using the Shannon diversity index, which was calculated using _SCIKIT_-_BIO_ v0.5.9 (Rideout et al., 2023). Diversity was estimated based on cellular organic carbon (C_OC_).

## 3 Results and discussion

### 3.1 Total coccolithophore relevance to global ocean carbon stocks

We developed a data-driven pipeline using an ensemble of machine-learning models (A_BIL_._PY_; (de Vries et al., 2025b, a)) to predict the abundance of the 58 most frequently occurring coccolithophore species from a set of environmental predictors. This pipeline was applied to interpolate the CASCADE coccolithophore abundance dataset (de Vries et al., 2024), using gridded monthly depth-revolved climatologies of environmental variables (see Methodology). We subsequently converted the predicted abundances into organic and inorganic carbon stocks using species-specific conversion factors derived from CASCADE (de Vries et al., 2024; Sheward et al., 2024). The data-driven models show good performance, with reasonable coefficient of determination (0.27 *±* 0.15 R^2^) and prediction errors (1.28 *±* 3.26 *µ*mol C m*^−^*^3^ Mean Average Error, and 4.90 *±* 15.16 *µ*mol C m*^−^*^3^ Root Mean Squared Error; data S1). Furthermore, the model predictions are primarily within the environmental range of the training dataset, with notable exceptions in regions with seasonal ice cover, and the North Pacific subpolar Ocean (Figure A4). The final outputs provide monthly, three-dimensional, global maps of abundance, biomass (cellular organic carbon, C_OC_), and calcium carbonate (cellular inorganic carbon, C_IC_) stock for each species.

Considering the entire coccolithophore community, our data-driven models estimate a global annual mean coccolithophore biomass of about 15.3 Tg C_OC_ [5.9, 29.1] integrated over the top 200 m of the ocean. Within the top 100 m, biomass is about 13.0 Tg C_OC_ [5.1, 23.4], which suggests that coccolithophores represent only a minor fraction of the total phytoplankton biomass, contributing about 2-5 % of the estimated range of 300-860 Tg C_OC_ for total phytoplankton in the photic zone ((Kostadinov et al., 2016) and references therein). Our coccolithophore biomass estimate is consistent with the previously reported MAREDAT range (1-30 Tg C_OC_ in the top 200 m (Buitenhuis et al., 2013)) but offers greater precision by representing spatiotemporal species-specific distributions.

For cellular inorganic carbon (C_IC_) our models estimate a global annual mean coccolithophore carbon stock of about 10.3 Tg C_IC_ [3.8, 20.1] in the upper 200 m and 8.4 Tg C_IC_ [3.1, 15.8] in the upper 100 m. This result represents the first depth- and monthly-resolved, species-specific estimate of the global coccolithophore C_IC_ stock. Our estimate is less than half of the Particulate Inorganic Carbon (PIC) values derived from satellite-based products (16.2-31.3 Tg PIC in the top 100 m) (Balch et al., 2005; Hopkins et al., 2019; Zhang et al., 2024). However, satellite-based PIC estimates include detached coccoliths (Balch and Mitchell, 2023; Neukermans et al., 2023). In contrast, our model focuses exclusively on C_IC_ while excluding detached coccoliths. The discrepancy between our estimate and satellite-derived estimates suggests that detrital inorganic carbon may play an important role in contributing to the total ocean PIC stocks, similar to how organic detrital material contributes to 51-86 % of ocean POC ((Kostadinov et al., 2016) and references therein).

Compared to a similar data-driven model estimate for other planktonic calcifiers, our findings indicate that global coccolithophore C_IC_ stock is about 50 times greater than planktonic foraminifera stock and *≈* 30 % lower than pteropod stock (0.2 and 14.1 Tg C_IC_ in the upper 200 m, respectively) (Knecht et al., 2023). Collectively, this results in a total living planktonic C_IC_ stock of about 24.6 Tg C_IC_, with coccolithophores accounting for about 42 %. This global comparison contrasts with direct observations in the North Pacific by Ziveri et al. (2023), and observations in the South Subtropical Atlantic by Kruijt et al. (2026) who both estimated that coccolithophores overall (combining living coccospheres and detached coccoliths) account for about 80 % of the total planktonic PIC stock in those regions. This discrepancy may be due to the contribution of detached coccoliths unaccounted for in our study and model-specific differences such as handling of pseudo-absences and unresolved depth and seasonal trends in Knecht et al. (2023)’s study. In addition, given that coccolithophore turnover rate is significantly faster than foraminifera and pteropods (about 1 d*^−^*^1^ relative to 0.1 and 0.05 d*^−^*^1^, respectively) (Ziveri et al., 2023), their contribution to C_IC_ production and export fluxes are amplified by at least an order of magnitude relative to their stock, leading to a significantly larger contribution to inorganic carbon fixation, as recently estimated (*≈* 90 %; (Ziveri et al., 2025)). These results highlight the critical role of coccolithophores in the marine carbon cycle, particularly in the context of ocean calcium carbonate cycling, reinforcing their importance in regulating the carbonate pump and ocean alkalinity.

### 3.2 A subtropical dominance of coccolithophore inorganic carbon stock

When considering super biomes (Figure 3D and Methods) depth-averaged coccolithophore C_IC_ concentrations are highest in subpolar regions (20.9 *µ*mol C m*^−^*^3^ [6.9, 44.1]; table 1), which contributes a major component to integrated stocks (29.6 % [8.6, 54.7]). This region includes the North Pacific, the Arctic Ocean, and the subpolar Southern Ocean and is characterized by large coccolithophore blooms (Balch et al., 2005; Hopkins et al., 2019). Subtropical Seasonally Stratified and Subtropical Permanently Stratified regions contain lower concentrations (14.4 *µ*mol C m*^−^*^3^ [4.6, 30.7] and 10.0 *µ*mol C m*^−^*^3^ [3.9, 17.7] respectively; table 1), but due to their large surface areas, contributions to integrated C_IC_ stocks of these biomes is comparable to the subpolar biomes (24.7 % [7.5, 48.7] and 35.4 % [14.1, 61.0] respectively; Figure 3E and table 1). When combining both subtropical biomes, this habitat becomes the largest contributor to coccolithophore C_IC_ stock (56.3 % [21.1, 83.1]; Figure A5). Concentrations in the Ice and Equatorial biomes are both low, and when integrated, are minor contributors to total C_IC_ stocks (Figure 3E and table 1). Our results match satellite PIC observations that *≈* 50 % of stock falls between 30° N and 30° S (Balch et al., 2005), with 40.6 % [13.1, 76.2] of C_IC_ stocks attributed to Equatorial and Subtropical Permanently Stratified biomes when combined.

**Figure 3.**
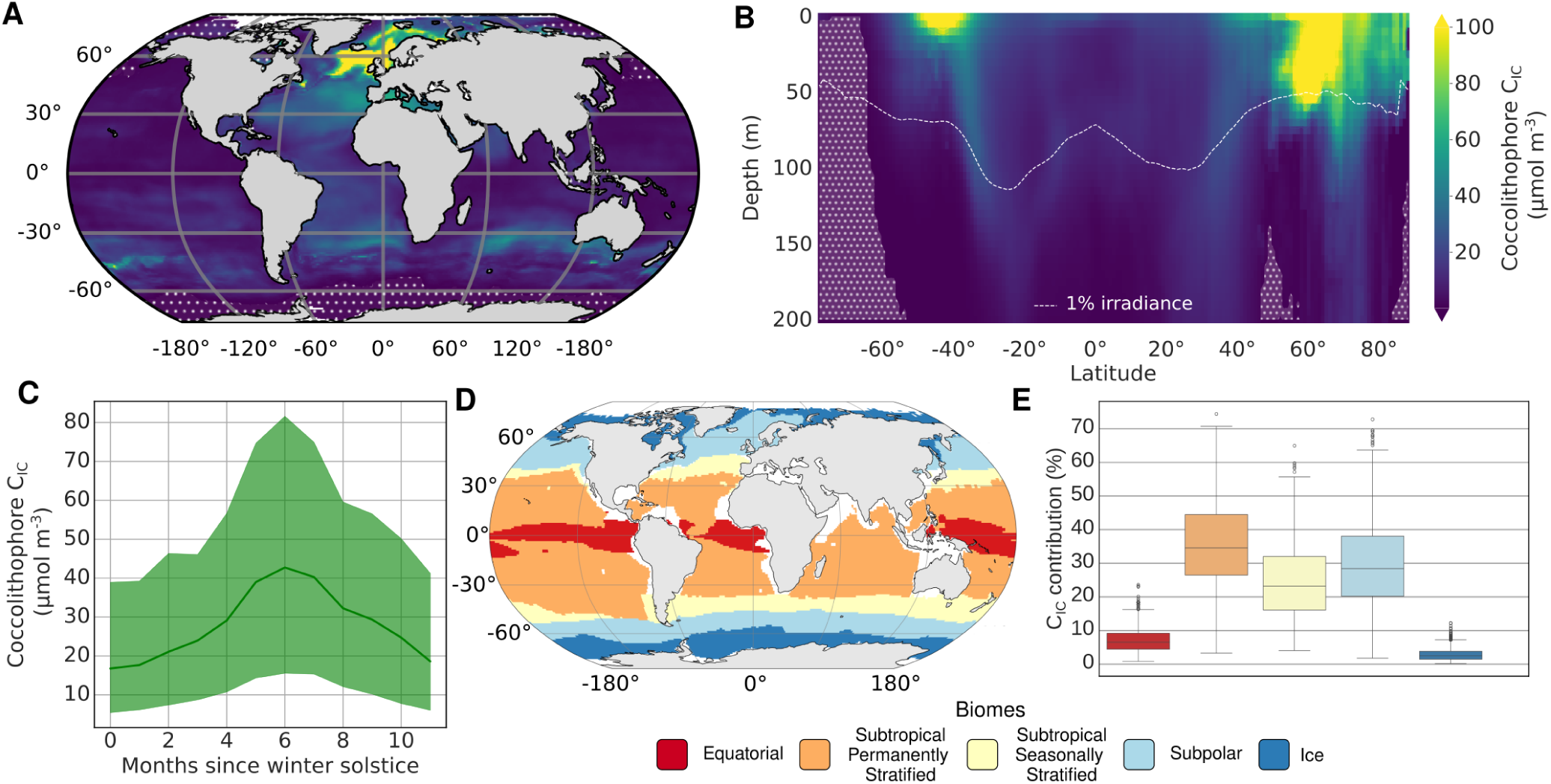
Spatial-temporal distributions of coccolithophore inorganic carbon stocks. **(A)** Depth and monthly averaged latitude-longitude concentration distributions. **(B)** Seasonally and longitudinally averaged latitude-depth concentration distributions. The 1 % surface irradiance threshold was derived from Castant et al., 2024 (Castant et al., 2024). **(C)** Temporal concentration distributions (shaded area denotes 95th percentile confidence intervals). **(D)** Superbiomes used in this study defined based on Fay and McKinley 2014 (Fay and McKinley, 2014). **(E)** Integrated stocks for each biome. Stippling in the map and latitude–depth section panels denotes regions where more than 50% of biomass predicted falls outside the training domain.

**Table 1.** C_IC_ concentrations and contributions to global stocks of the different biomes. Brackets denote 95th percentile confidence intervals.

| biome | concentration<br>($\mu\text{mol C m}^{-3}$) | contribution<br>(%) |
| --- | --- | --- |
| Global | 11.3 [4.1, 21.8] | NA |
| Ice | 7.6 [2.1, 15.4] | 2.9 [0.5, 7.5] |
| Subpolar | 20.9 [6.9, 44.1] | 29.62 [8.6, 54.7] |
| Subtropical Seasonally Stratified | 14.36 [4.6, 30.7] | 24.7 [7.5, 48.7] |
| Subtropical Permanently Stratified | 10.0 [3.9, 17.7] | 35.4 [14.1, 61.0] |
| Equatorial | 5.6 [2.0, 10.8] | 7.4 [1.9, 16.2] |

Seasonally, the total coccolithophore C_IC_ concentrations present a summer bloom (Boreal July and Austral January) at 42.7 *µ*mol C m*^−^*^3^ [15.5, 81.6] which broadly matches satellite PIC phenology (Hopkins et al., 2015). Seasonality becomes more strongly pronounced with increasing latitude, with the highest fluctuation in subpolar biomes (Figure 4). This variable phenology further magnifies the net contributions of subtropical biomes to global C_IC_ stocks.

**Figure 4.**
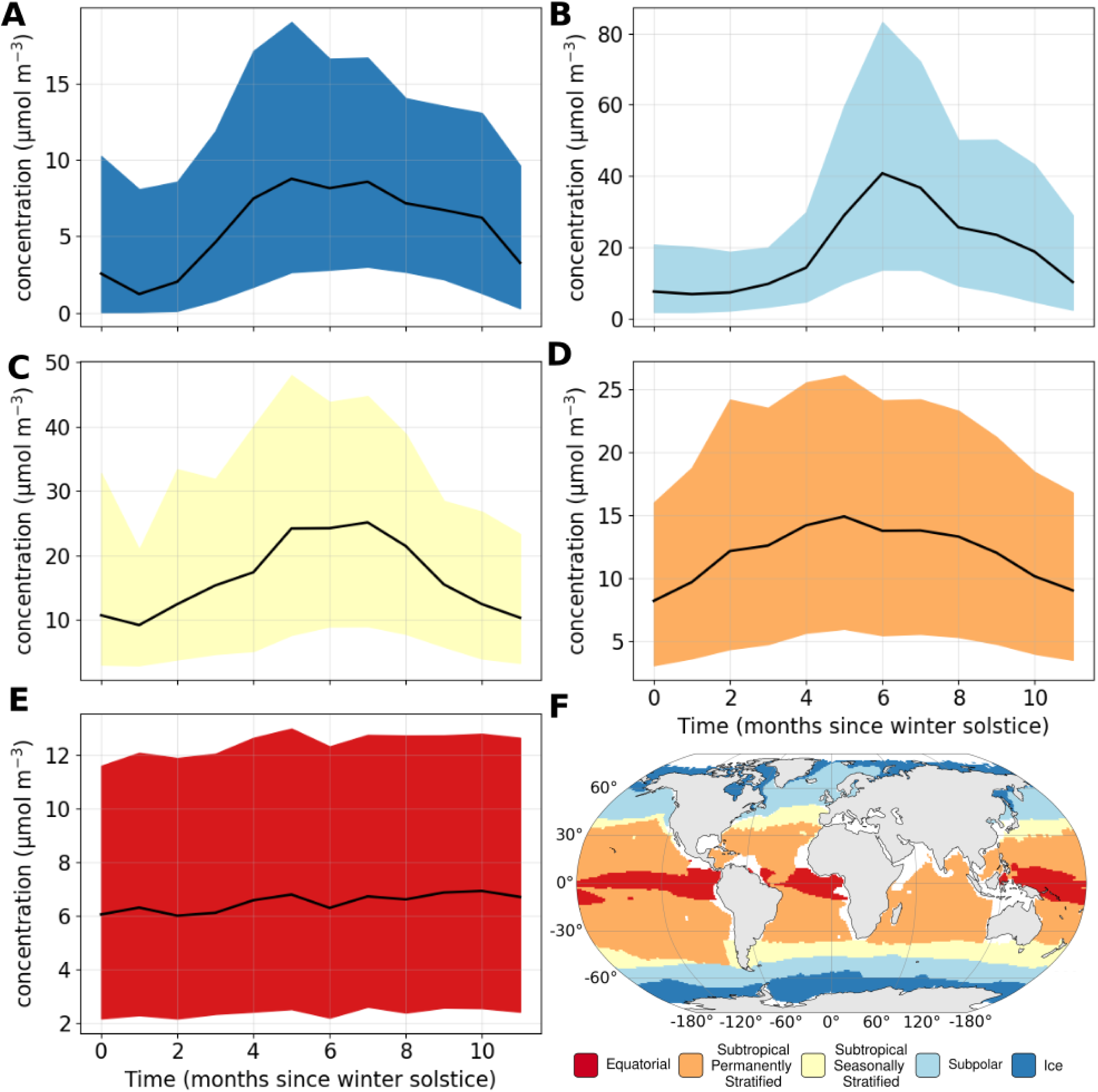
Seasonality of Cellular Inorganic Carbon (C_IC_) concentrations of the different biomes. **(A)** Ice. **(B)** Subpolar. **(C)** Equatorial. **(D)** Subtropical Seasonally Stratified. **(E)** Subtropical Permanently Stratified. **(F)** Equatorial. **(G)** Spatial distribution of biomes. Biomes from Fay and McKinley 2014 (Fay and McKinley, 2014). Shaded areas denote 95th percentile confidence intervals.

Thus, while our model illustrates the high concentrations of coccolithophore C_IC_ in subpolar regions, our results also highlight the importance of the subtropics, and a need to consider surface area and seasonality when determining global coccolithophore C_IC_ stocks.

### 3.3 A third of inorganic carbon stock is found below the Euphotic Zone

Vertically, our models reveal a distinct latitudinal trend in coccolithophore C_IC_ stock (Figure 3B). In the subpolar regions, the C_IC_ stock presents a typical exponential decline with depth, maintaining high concentrations (*>* 70 *µ*mol C m*^−^*^3^) within the upper 25 m in the Southern Ocean and up to 75 m in the subpolar North Atlantic. Conversely, inorganic carbon in the subtropical region is uniformly distributed with depth, maintaining medium concentrations (*<* 20 *µ*mol C m*^−^*^3^) in the Sub-Euphotic Zone (*<* 1 % of surface irradiance levels) (Poulton et al., 2017), Figure 3B, dashed line). When considering integrated stock for the Upper Euphotic Zone (*>* 10 % surface irradiance), the Lower Euphotic Zone (1-10 %) and the Sub-Euphotic Zone (as defined in Poulton et al. (2017)), relative contributions to global C_IC_ are 41.0 % [17.1, 68.0], 27.1 % [8.3, 52.4] and 32.0 % [9.8, 59.4] respectively (Figure A6).

Relatively high stocks within the Sub-Euphotic Zone have been observed previously (Poulton et al., 2017). While sinking could explain some of this stock, species inhabiting this zone have unique communities and are highly specialized (Poulton et al., 2017). Furthermore, sinking cells might nonetheless be alive (Agusti et al., 2015). Finally, many coccolithophores are expected to be phagotrophic (Poulton et al., 2017; Gibbs et al., 2020) and/or osmotrophic (Flynn et al., 2013), potentially offsetting light limitation. Nonetheless, osmotrophic uptake and predation rates vary greatly between species (Ziveri et al., 2025) raising open questions around net production rates in the Sub-Euphotic Zone.

### 3.4 *Gephyrocapsa huxleyi* contributes less than 10 percent to global C_IC_ stock

In agreement with previous studies (de Vries et al., 2021), our global model demonstrates that *G. huxleyi* dominates total annual mean coccolithophore abundance, contributing 28.5 % [12.3, 46.4] (Table A2) of the total (4.8 x 10^23^ cells [1.9 x 10^23^, 8.9 x 10^23^]). Although this contribution is lower than estimated previously (59 %) (de Vries et al., 2021), the previous estimate relied on taking the mean from an abundance dataset and is thus skewed by observed sampling effort biases towards the surface, summer and subpolar regions where *G. huxleyi* abundances are highest.

In contrast to cellular abundances, our models estimate that *G. huxleyi* contributes to less than ten percent of the total annual mean coccolithophore C_IC_ stock, accounting for 7.2 % [2.4, 14.7] (Figure 5B, Table A4). This relatively small contribution is due to *G. huxleyi* lower cellular calcite content and C_IC_:C_OC_ ratio compared to other abundant coccolithophore species (Figure 1). Regionally, *G. huxleyi* C_IC_ stock is highest in subpolar biomes where it contributes 16.8 % [4.8, 37.7] of integrated C_IC_ stocks (Table A3). Seasonally, *G. huxleyi* C_IC_ concentrations peak in the late summer months (Figure 5M).

**Figure 5.**
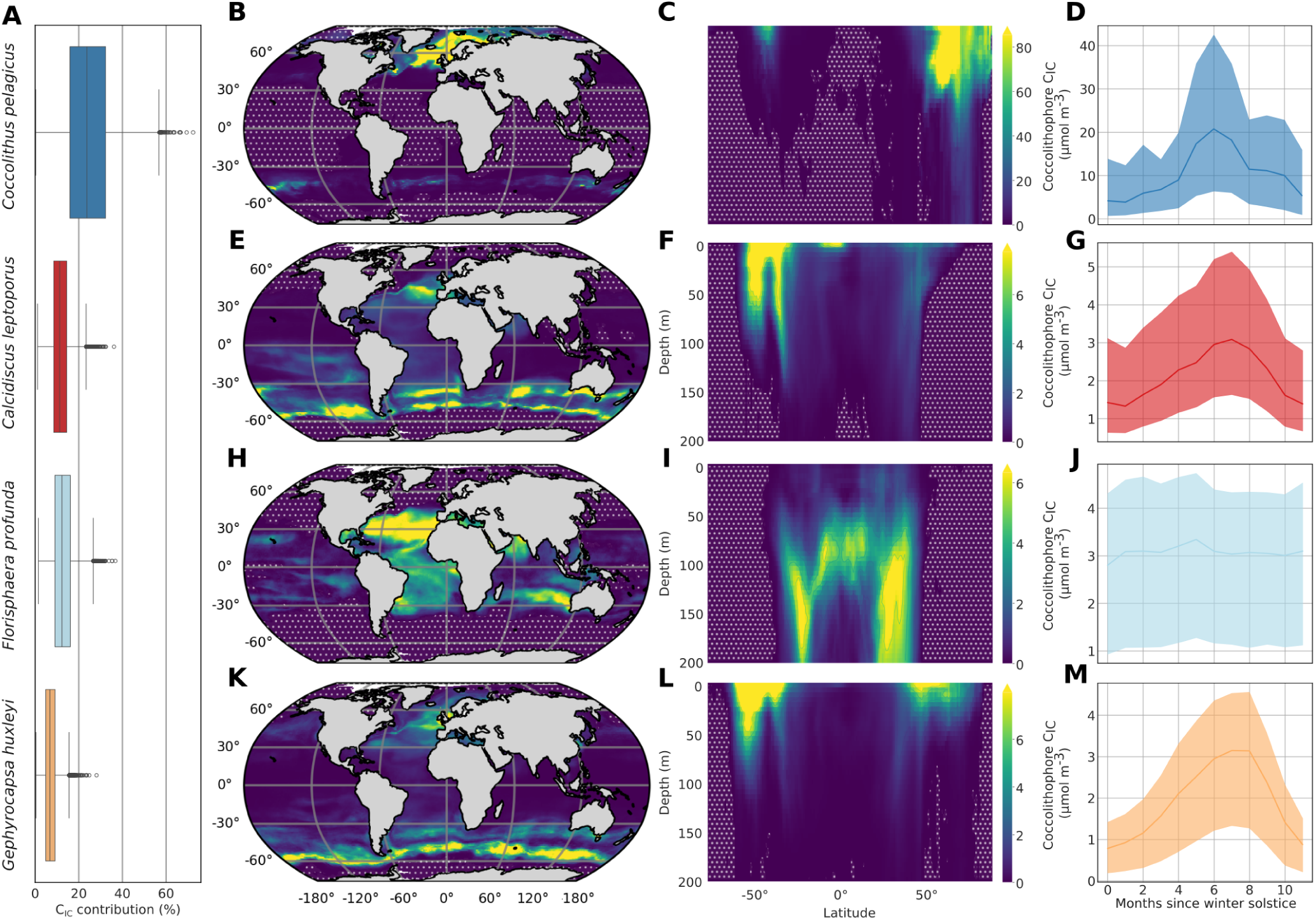
Spatio-temporal distributions of primary coccolithophore contributors to inorganic carbon stocks (C_IC_), including *Coccolithus pelagicus*, *Calcidicus leptoporus*, *Florisphaera profunda* and *Gephyrocapsa huxleyi*. (**A**) Relative contributions of the top species to total C_IC_ stock (%). **(B, E, H, K)** Latitude-longitude distributions of individual species averaged with depth and month (*µ*mol C_IC_ m*^−^*^3^). **(C, F, I, L)** Latitude-depth distributions of individual species averaged with depth and month (*µ*mol C_IC_ m*^−^*^3^). **(D, G, J, M)** Seasonal distributions of individual species averaged with latitude, longitude and depth (*µ*mol C_IC_ m*^−^*^3^). Seasonal plots denote the number of months since the winter solstice.

Thus while *G. huxleyi* is a key contributor to C_IC_ stocks, our models illustrate that a significant proportion of coccolithophore stocks is attributed to other less studied species.

### 3.5 Important contribution of large heavily calcified coccolithophore species

Our model shows that in addition to *G. huxleyi*, large and heavily calcified species contribute majorly to global inorganic carbon stocks. These species include large contributions of *Coccolithus pelagicus* (24.7 % [5.7, 48.0]), *Calcidicus leptoporus* (11.7 % [4.7, 21.7]) and *Florisphaera profunda* (13.0 % [4.7, 24.6]) (Table A3). Although these species have low relative abundances (Table A2), their large contributions are driven by their larger cell size and elevated cellular C_IC_: C_OC_ ratio (Figure 1). Collectively, these three species account for around half of the total C_IC_ stock. These findings show the disproportionate influence of a small number of species, particularly heavily calcifying ones, on coccolithophore driven carbon cycling, with just three out of approximately 250-300 known (morpho)species responsible for half of the global coccolithophore carbonate stock.

In our model, *C. leptoporus* is most abundant in the subtropical region of the North Atlantic and the subtropical and subpolar regions of Southern Ocean and consequently is a key contributor to both the subpolar and subtropical seasonally stratified biome C_IC_ stocks (12.9 % [3.3, 31.0] and 20.2 % [7.5, 36.7], respectively) (Table A3). In contrast, *C. pelagicus* coccolithophore C_IC_ stock is highest in the Subpolar North Atlantic region and the Subtropical Seasonally Stratified Southern Ocean, consequently contributing majorly to both the Subpolar (64.3 % [29.5, 86.3]) and Subtropical Seasonally Stratified biomes (30.8 % [3.7, 63.3]) (Table A3). Finally, *F. profunda* C_IC_ stock is highest in the Equatorial and Subtropical Permanently Stratified biomes where it constitutes 21.5 % [7.9, 38.7] and 20.5 % [8.6, 34.9] of C_IC_ stock respectively (Table A3).

Like *G. huxleyi*, *C. leptoporus* and *C. pelagicus* preferentially grow in surface waters, with highest concentrations within the first 50 m (Figure 5D, H). In contrast, *F. profunda* is a deep-dwelling coccolithophore species that preferentially grows in the Sub-Euphotic Zone (*<* 1 % surface irradiance)(Poulton et al., 2017) and shows peak concentrations of 6 *µ*mol C m*^−^*^3^ in this region (Figure 5L). Furthermore, this species shows low seasonality in contrast to the other dominant species (Figure 5M).

Furthermore, *C. leptoporus* and *C. pelagicus* are non-motile and form well-defined circular shape coccoliths, which are termed placoliths and which cover the entire cell (Young, 1994). In contrast, *F. profunda* is covered by nanoliths, is motile, and features a large coccosphere opening (Young, 1994). While all three species have chloroplasts (Young et al., 2022), and can thus undergo photosynthesis, the deep dwelling nature of *F. profunda* suggests that this species might be mixotrophic (Poulton et al., 2017). However, unlike the other three dominant coccolithophores, *F. profunda* has not yet been successfully cultured in the laboratory, limiting our understanding of its biology and ecological function.

### 3.6 Uneven contribution of a diverse coccolithophore community

In addition to *G. huxleyi*, only three other species, *F. profunda*, *C. leptoporus* and *C. pelagicus*, contribute more than 5 % to global C_IC_ stock of the 58 species included in our model (Table A4). Nonetheless, rare species (*≤* 5 % contribution) constitute 43.3 % [28.7, 58.6] of global C_IC_ stocks (Table A3). Furthermore, cumulative summations (see Methods) suggest that 13 species are needed to reach *>* 80 % of global C_IC_ stocks, while 21 species are needed to reach *>* 80 % of global C_OC_ stocks. Contributions of such rare species are particularly high in the Subtropical Permanently Stratified and Equatorial regions (61.4 % [48.5, 73.9], 68.5 % [52.3, 82.8], respectively; Table A3). Our study thus highlights the importance of rare coccolithophore species to carbon stocks, particularly in the low latitudes.

The distribution of rare species is reflected in the Shannon diversity index, with the highest diversity observed in Subtropical biomes and lowest diversity observed in Subpolar and Ice biomes (Figure 6A-B, see Methods). Notably, diversity is also lower in equatorial upwelling zones compared to subtropical regions. This pattern is consistent with previous studies that characterized coccolithophore diversity (O’Brien et al., 2016; Benedetti et al., 2023) and generally matches other planktonic organisms (Benedetti et al., 2023). More broadly, high biodiversity in the (sub)tropics is observed across all kingdoms of life, pointing towards a universal mechanism (Tittensor et al., 2010). While mechanisms of latitudinal diversity gradients are of debate (Tittensor et al., 2010), our vertical diversity trends which show a decrease in diversity with depth in combination with latitudinal patterns illustrate that low diversity is in part driven by the competitive exclusion of *G. huxleyi*, *C. leptoporus*, *C. pelagicus* and *F. profunda* in regions they dominate.

**Figure 6.**
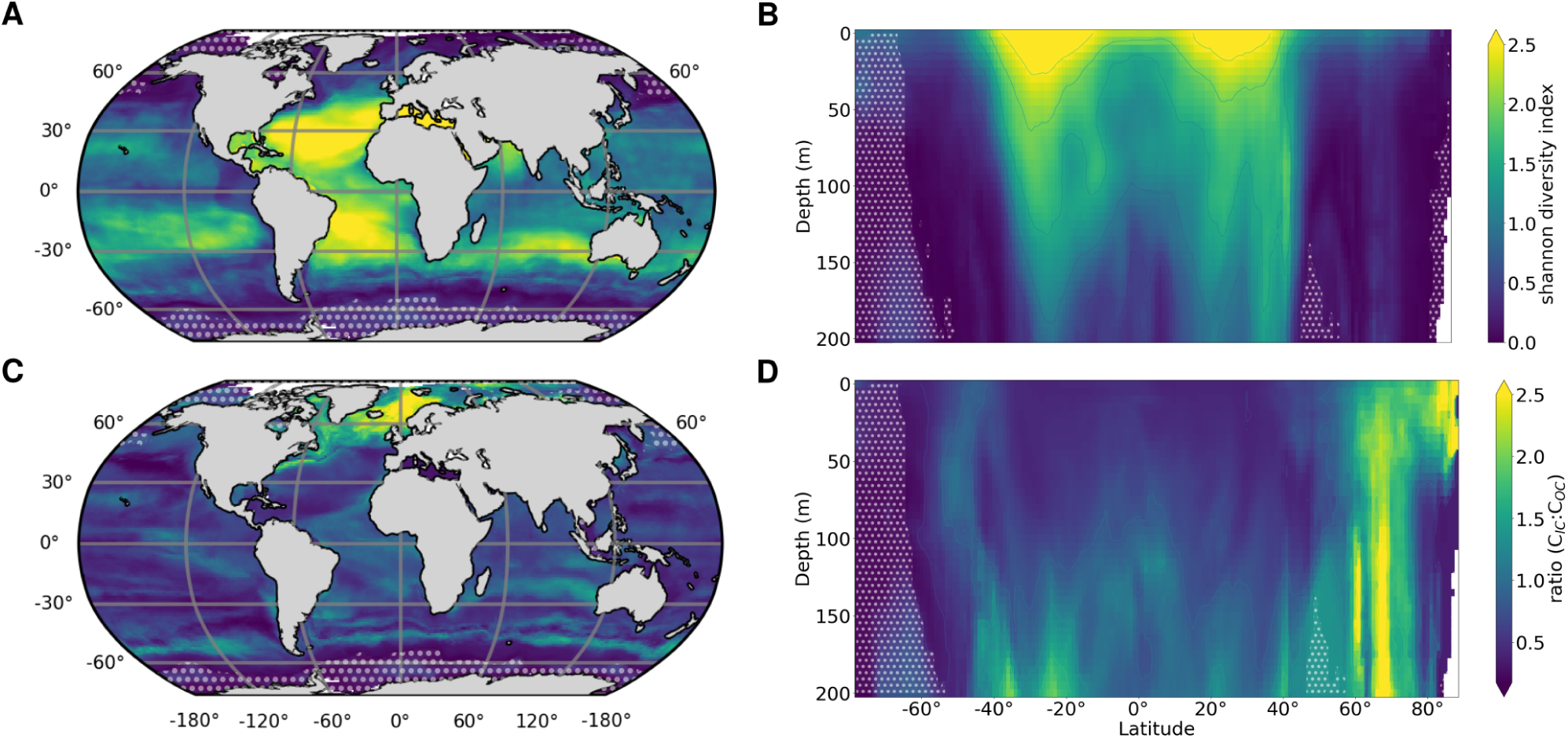
Spatial distributions of coccolithophore diversity and cellular inorganic to organic carbon ratios. **(A)** Latitude-longitude distributions of Shannon diversity. **(B)** Depth-latitude distributions of Shannon diversity. **(C)** Latitude-longitude distributions of cellular organic and inorganic carbon (C_IC_:C_OC_) contents. **(D)** Latitude-depth distributions of C_IC_:C_OC_.

### 3.7 Decoupled coccolithophore inorganic and organic carbon stocks

The difference in distributions among the top coccolithophore carbonate contributors and their cellular calcite content in our models means that, on a global scale, total coccolithophore C_IC_ and C_OC_ distributions are decoupled (Figure 6A-B, Figure A7). Specifically, the global distribution of total coccolithophore C_IC_:C_OC_ ratio is not uniform, presenting distinct latitudinal and vertical trends. Elevated C_IC_:C_OC_ ratios, around 1.5-2.2, are observed in deep subtropical regions and the subpolar North Atlantic and Arctic, due to the dominant presence of the heavily calcified *Florisphaera profunda* in the subtropics and *Coccolithus pelagicus* in subpolar regions. In contrast, lower C_IC_: C_OC_ ratios (*<* 1.5) are found elsewhere.

This spatial heterogeneity in the total coccolithophore C_IC_:C_OC_ ratio has important implications for climate model representations of calcification. The few models in the Climate Model Intercomparison Project (CMIP) that explicitly include calcification assume a fixed ratio between inorganic and organic carbon production ((Planchat et al., 2023), and references within). Not accounting for C_IC_:C_OC_ spatial heterogeneity may misrepresent interactions with other spatially varying processes such outgassing and export, thus potentially leading to mischaracterizations of such processes. Since spatial heterogeneity of C_IC_:C_OC_ is driven by species-specific distribution patterns, our results suggest that CMIP model representation of calcium carbonate distributions could be improved by explicitly representing variable C_IC_:C_OC_ ratios, as expressed by different coccolithophore plankton functional types (PFTs). While significant progress has already been made to incorporate coccolithophore PFTs into climate models (Le Quéré et al., 2016; Seifert et al., 2022; Krumhardt et al., 2019), these models have been primarily parameterized by the small and lightly calcified coccolithophore species *G. huxleyi*. Since important model parameterizations are influenced by cell size ((Litchman and Klausmeier, 2008) and reference therein), C_IC_:C_OC_ ratio (Gafar et al., 2019; Kottmeier et al., 2022), mixotrophy and motility (Ziveri et al., 2025) such models are unable to represent the remainder of the coccolithophore community constituting the majority of C_IC_ stocks. Furthermore, while traits like size and C_IC_:C_OC_ ratio are well understood, other traits like motility, mixotrophy and the benefit of calcification are poorly constrained (Monteiro et al., 2016) highlighting key challenges in modelling the ocean carbonate cycle.

### 3.8 Model limitations and outlook

Our results are reported with 95% confidence intervals (CIs), which can span a broad range for some estimates. Such uncertainty is expected given the sparse and patchy nature of plankton observations. In addition, these intervals are not directly comparable to the standard deviations (SDs) reported for other calcifying groups (Knecht et al., 2023), because SDs and CIs summarize uncertainty differently. When converted to approximate coefficients of variation, our integrated stock estimates show lower variability than reported for other calcifying groups (Knecht et al., 2023). Nonetheless, uncertainty could conceivably be reduced through additional sampling efforts, particularly in the North Pacific and sub-polar Sub-Euphotic zone as illustrated by our area of applicability analysis (Figure A4).

Aside from spatio-temporal biases, our study focuses on species-level contributions, ignoring sub-species variation in organic and inorganic carbon contents due to limited data availability in existing data compilations (de Vries et al., 2024; Sheward et al., 2024). Further efforts to estimate sub-species variation in these conversion factors would thus likewise be a worthwhile endeavour.

Finally, our study only considers stocks, and does not estimate carbon fixation rates which is a function of both stocks and per-capita growth rates. Since growth rates can vary between species, largely driven by size (Villiot et al., 2021) this is will be an important step for better constraining species-specific contributions to the calcium carbonate cycle. Nonetheless, previous work suggests that even for species with different maximum growth rates, similar growth rates are observed under similar environmental conditions (Daniels et al., 2014).

## 4 Conclusion

This study provides the first species-specific quantification of the global coccolithophore CaCO_3_ stock. We find an important role of the Subtropical biomes which constitute the majority of inorganic carbon stocks, and the Sub-Euphotic Zone, where approximately a third of the stock is found. While *Gephyrocapsa huxleyi* is often assumed to be the primary contributor, our data-driven model shows that it accounts for less than 10 % of the total stock. Instead, 3 heavily calcified species (*Florisphaera profunda*, *Calcidicus leptoporus* and *Coccolithus pelagicus*) emerge as major contributors, collectively accounting for half of the global coccolithophore CaCO_3_ stock. Furthermore, 13 species are needed to constrain 80 % of coccolithophore stocks, illustrating that a diverse community constitutes the coccolithophore inorganic carbon budget.

Our results highlight that the coccolithophores that constitute the majority of coccolithophore stocks vary widely in their traits that determine their response to climate change such as size, C_IC_: C_OC_ ratio, motility, and mixotrophy. As such, coccolithophore response to climate change cannot be approximated by a single species. Our results thus highlight expanding the representation of non-*G. huxleyi* species in future lab characterizations and modelling efforts, is a prerequisite to accurately assessing the role of coccolithophores in ocean carbon cycling.

## Code and data availability

Model outputs from this study can be found on Zenodo (Abundance: https://doi.org/10.5281/zenodo.16886603; C_OC_: https://doi.org/10.5281/zenodo.16986899; C_IC_: https://doi.org/10.5281/zenodo.16887386).

The abundance and cellular carbon dataset used to train our model was obtained from the CASCADE dataset (de Vries et al., 2024), and is available at https://doi.org/10.5281/zenodo.13919902.

The software used to generate our results was obtained from the Abil.py GitHub repository (de Vries et al., 2025b (de Vries et al., 2025a), v25.08.06: https://doi.org/10.5281/zenodo.16886568).

The environmental data used for predictions is available on Zenodo: https://doi.org/10.5281/zenodo.16753617.

## Appendix A

**Figure A1.**
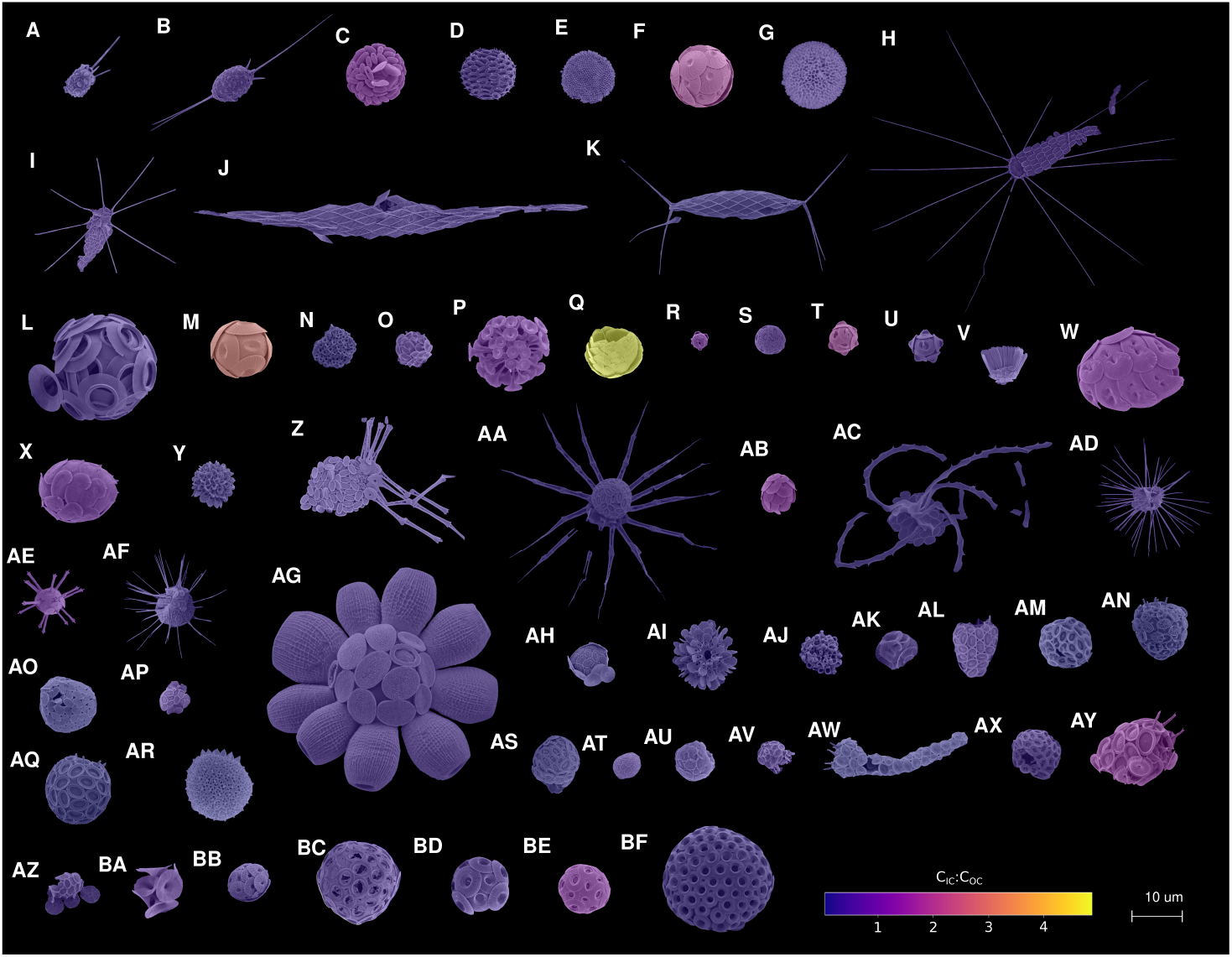
Morphological diversity of the coccolithophore species considered in this study. **(A)** *Acanthoica acanthifera*. **(B)** *Acanthoica quattrospina*. **(C)** *Algirosphaera robusta*. **(D)** *Alisphaera unicornis*. **(E)** *Alisphaera unicornis* POL. **F)** *Calcidiscus leptoporus*. **(G)** *Calcidiscus leptoporus* HOL. **(H)** *Calciopappus caudatus*. **(I)** *Calciopappus rigidus*. **(J)** *Calciosolenia brasiliensis*. **(K)** *Calciosolenia murrayi*. **(L)** *Ceratolithus cristatus*. **(M)** *Coccolithus pelagicus*. **(N)** *Corisphaera gracilis*. **(O)** *Cyrtosphaera aculeata*. **(P)** *Discosphaera tubifera*. **(Q)** *Florisphaera profunda*. **(R)** *Gephyrocapsa ericsonii*. **(S)** *Gephyrocapsa huxleyi*. **(T)** *Gephyrocapsa muellerae*. **(U)** *Gephyrocapsa oceanica*. **(V)** *Gladiolithus flabellatus*. **(W)** *Helicosphaera carteri*. **(X)** *Helicosphaera hyalina*. **(Y)** *Helladosphaera cornifera*. **(Z)** *Michaelsarsia adriaticus*. **(AA)** *Michaelsarsia elegans*. **(AB)** *Oolithotus antillarum*. **(AC)** *Ophiaster hydroideus*. **(AD)** *Palusphaera vandelii*. **(AE)** *Rhabdosphaera clavigera*. **(AF)** *Rhabdosphaera xiphos*. **(AG)** *Scyphosphaera apsteinii*. **(AH)** *Syracosphaera anthos*. **(AI)** *Syracosphaera anthos* HOL. **(AJ)** *Syracosphaera arethusae* HOL. **(AK)** *Syracosphaera corolla*. **(AL)** *Syracosphaera dilatata*. **(AM)** *Syracosphaera halldalii*. **(AN)** *Syracosphaera histrica*. **(AO)** *Syracosphaera histrica* HOL. **(AP)** *Syracosphaera marginiporata*. **(AQ)** *Syracosphaera mediterranea*. **(AR)** *Syracosphaera mediterranea* HOL wettsteinii type. **(AS)** *Syracosphaera molischii*. **(AT)** *Syracosphaera nana*. **(AU)** *Syracosphaera nodosa*. **(AV)** *Syracosphaera ossa*. **(AW)** *Syracosphaera prolongata*. **(AX)** *Syracosphaera protrudens*. **(AY)** *Syracosphaera pulchra*. **(AZ)** *Syracosphaera rotula*. **(BA)** *Umbellosphaera irregularis*. **(BB)** *Umbellosphaera tenuis*. **(BC)** *Umbilicosphaera anulus*. **(BD)** *Umbilicosphaera foliosa*. **(BE)** *Umbilicosphaera hulburtiana*. **(BF)** *Umbilicosphaera sibogae*. All images by Jeremy Young except *Acanthoica acanthifera* by Annelies Kleijne, *Rhabdosphaera xiphos* by Luka Šupraha and *Syracosphaera halldalii* by Elisa Malinverno. Images were acquired from Nannotax3 (Young et al., 2022). The C_IC_:C_OC_ data was acquired from (de Vries et al., 2024).

**Figure A2.**
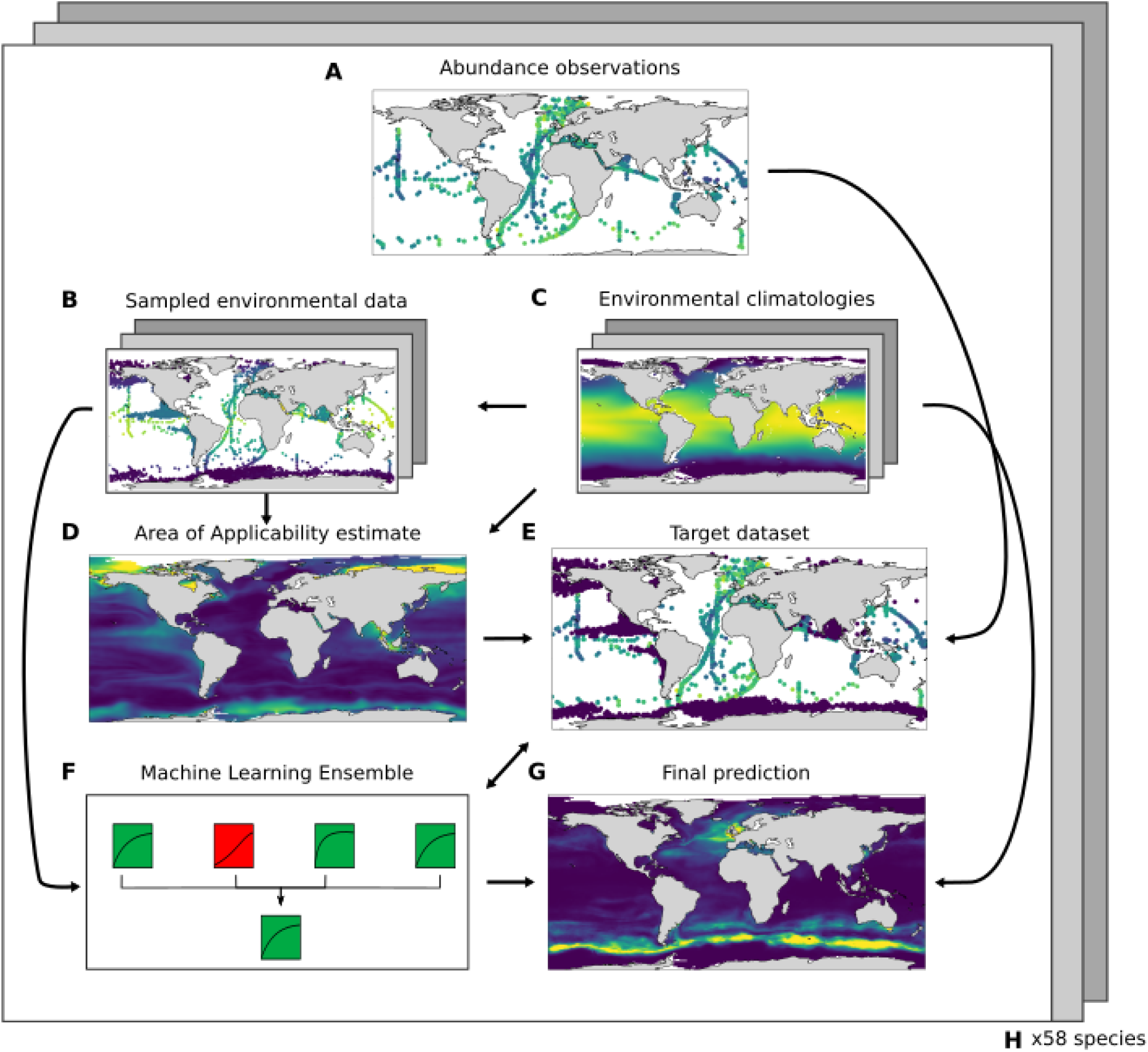
Machine Learning pipeline used in this study. **(A)** Biomass observations are derived from CASCADE (de Vries et al., 2024) for each species. **(B)** Relevant environmental data is sampled for each biomass observations from **(C)** climatologies of environmental data. **(D)** Area of Applicability (AOA) is estimated based on sampled environmental data and the full environmental climatologies. **(E)** Target data to be predicted by the Machine Learning (ML) pipeline is generated by introducing pseudo absences to the biomass observations based on the AOA estimates. **(F)** A ML ensemble is trained using 10-fold cross validation. **(G)** Interpolated prediction are made based on the climatology and ML ensemble. **(H)** The process is repeated for each species.

**Figure A3.**
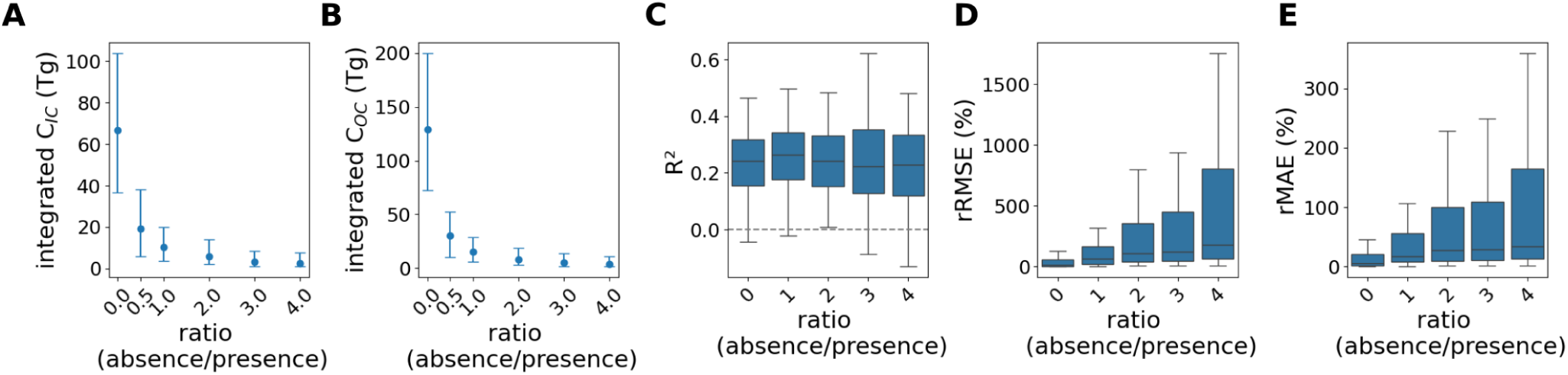
Integrated stock and error metrics of simulations with different pseudo absence to presence ratios. **(A)** Integrated cellular inorganic carbon (C_IC_). **(B)** integrated cellular organic carbon (C_OC_). **(C)** Coefficients of determination (R^2^). **(D)** Relative Root Mean Squared Error (rRMSE). **(E)** Relative Mean Average Error (rMAE). Note that for this study a ratio of 1:1 was used.

**Figure A4.**
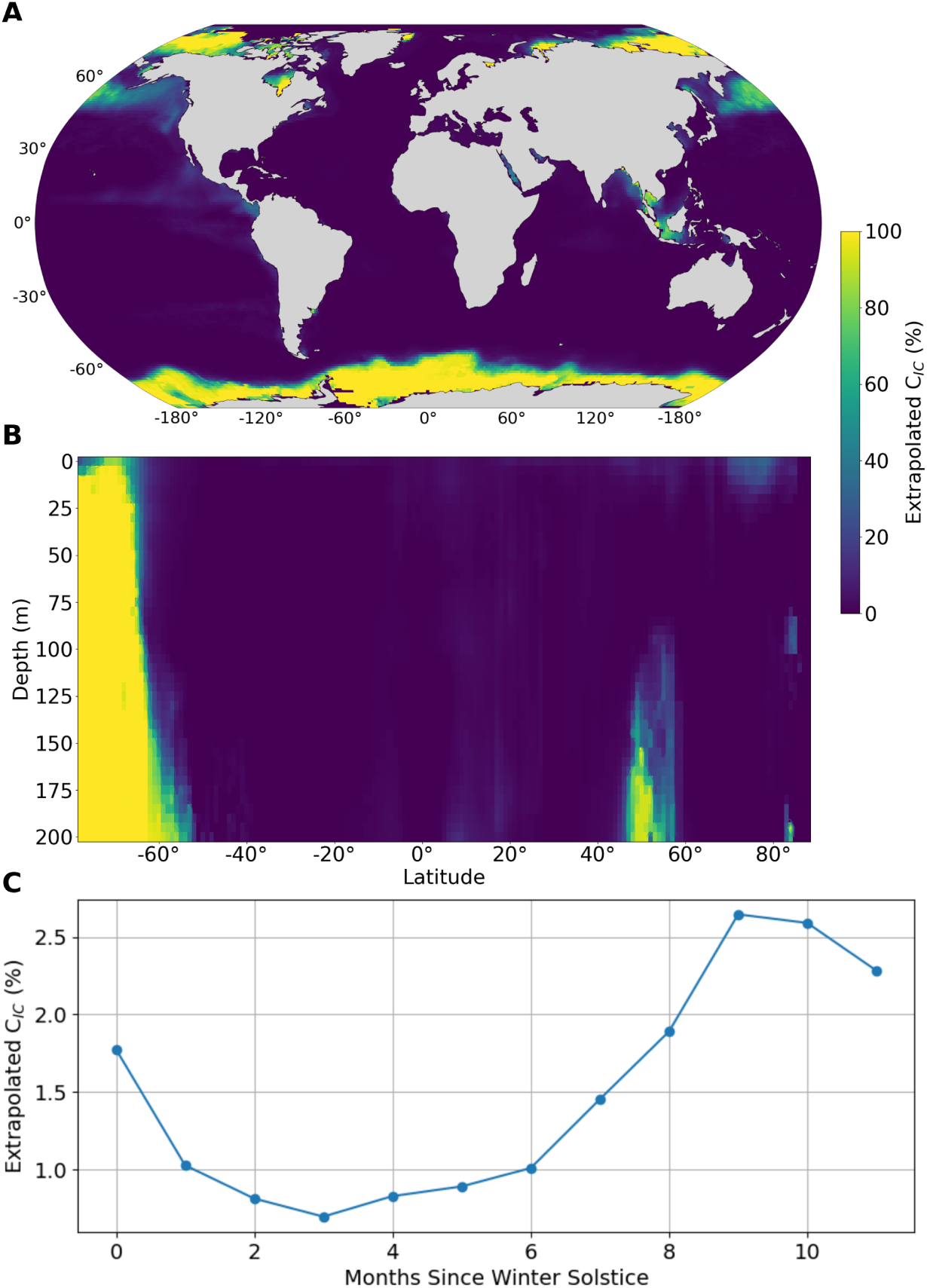
Spatiotemporal distributions of model cellular inorganic carbon (C_IC_) extrapolation. **(A)** Latitude-longitude extrapolation. **(B)** Latitude-depth extrapolation. **(C)** Temporal extrapolation. Note that for the temporal extrapolation, months since winter solstice was used to account for hemispherical phenology differences.

**Figure A5.**
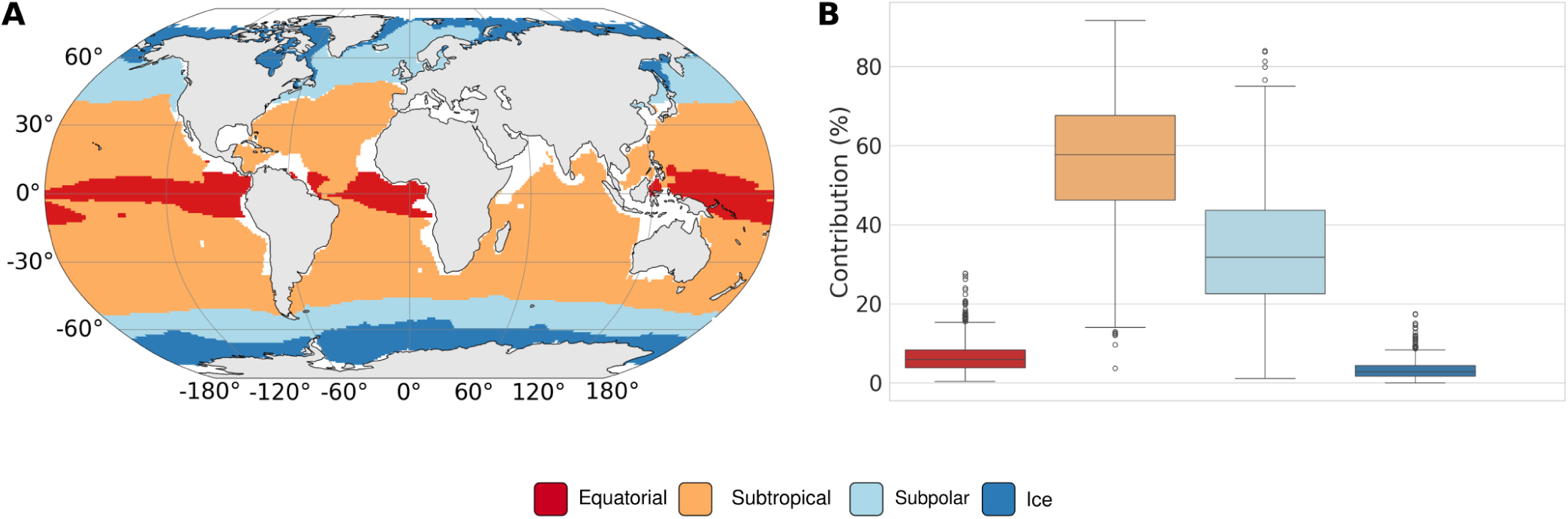
Contributions of biomes when subtropical biomes are merged. **(A)** Merged biomes, which combine the Subtropical Seasonally Stratified and Subtropical Permanently Stratified biomes from Fay and McKinley 2014 (Fay and McKinley, 2014). **(B)** Relative contributions to C_IC_ of each biome.

**Figure A6.**
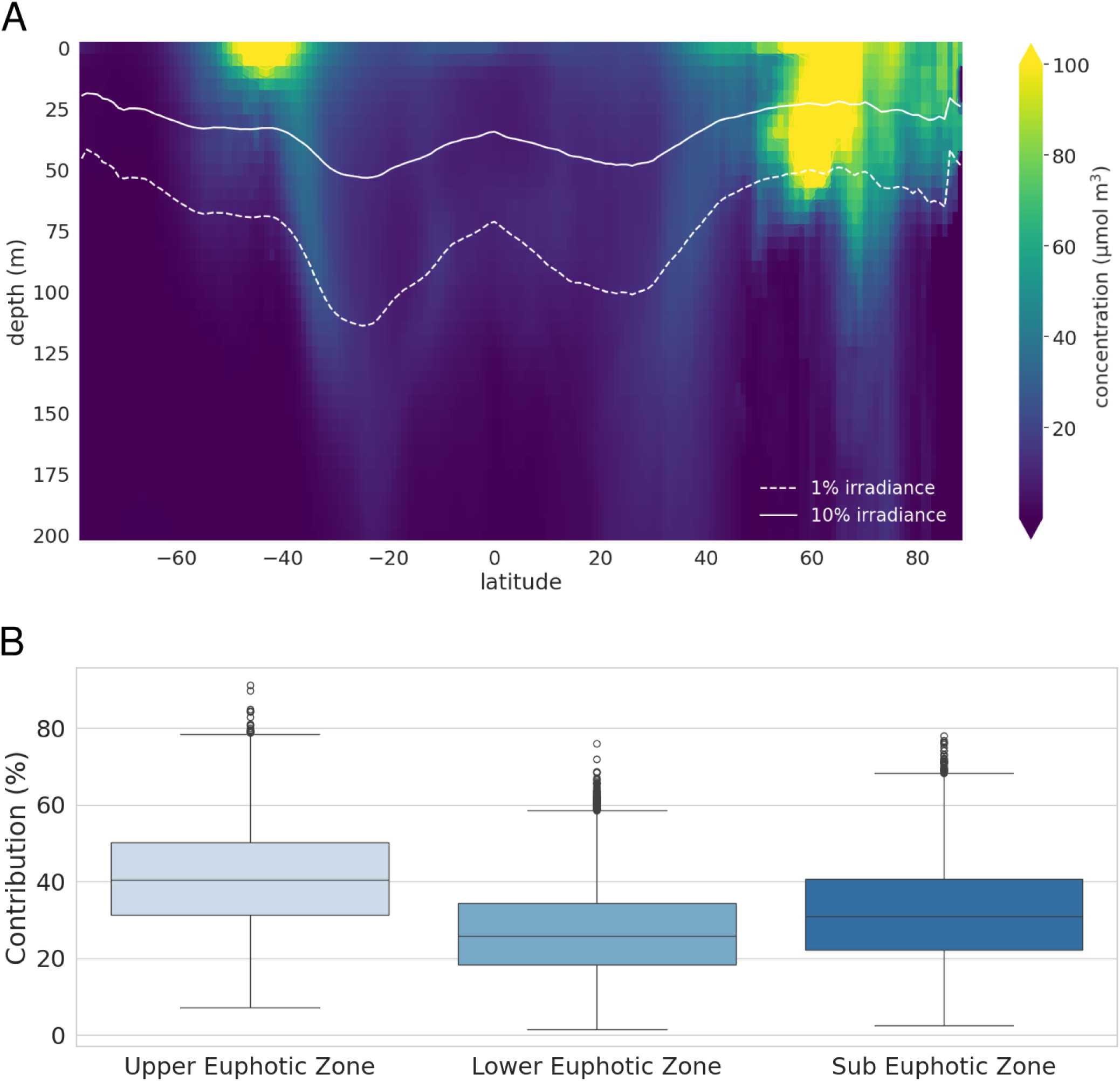
Photic zones and their relative contributions to integrated cellular inorganic carbon (C_IC_) stocks. **(A)** The zones are split into three categories: Upper photic zone (10-100 % of surface irradiance, Lower photic zone (1-10 %), and Sub-Euphotic zone (*<*1 %). **(B)** Resampled contributions for each photic zone.

**Figure A7.**
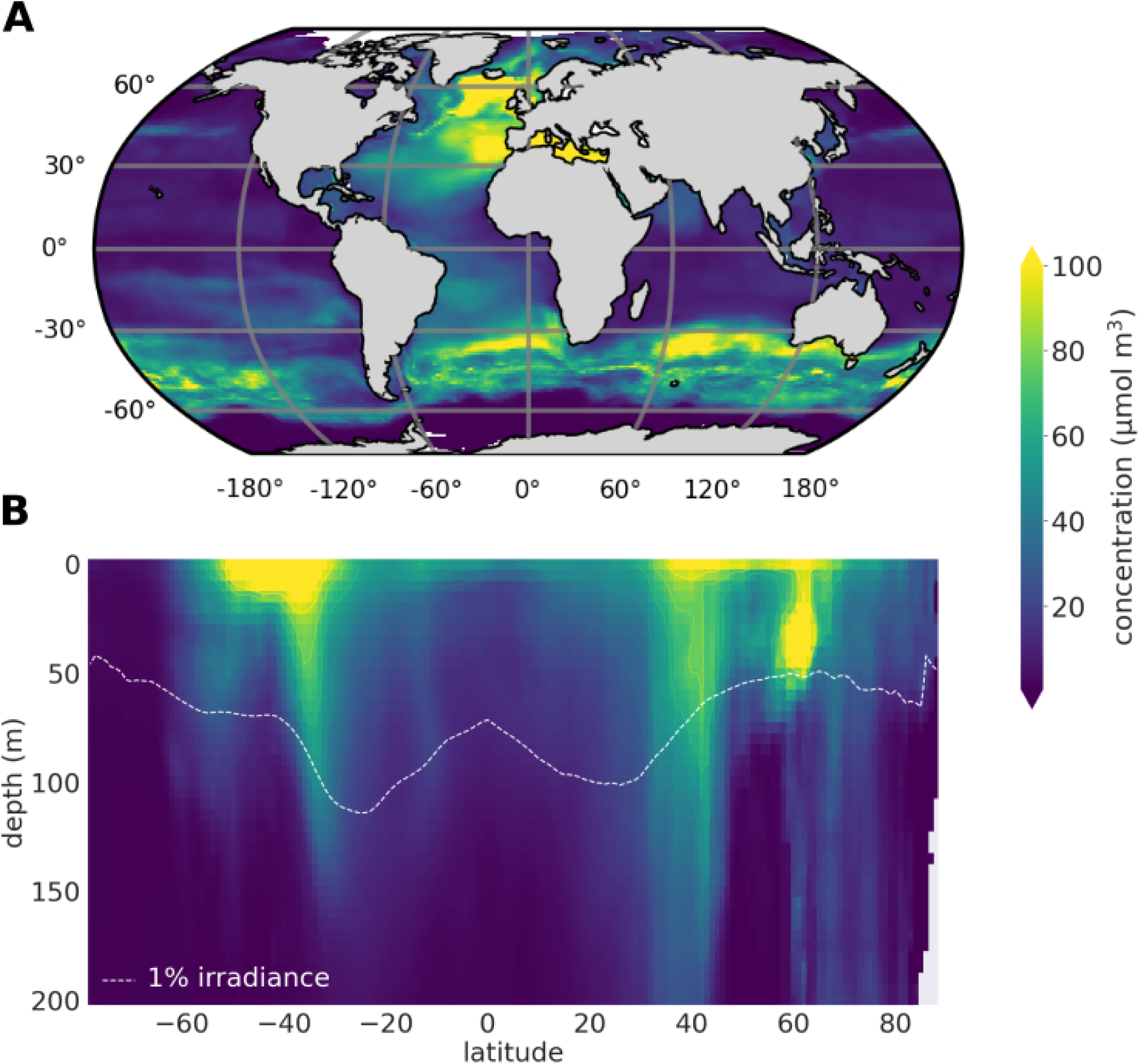
Spatial-temporal distributions of cellular organic carbon (C_OC_) stocks. **(A)** Depth and season averaged latitude-longitude concentration distributions. **(B)** Seasonally and longitudinally averaged latitude-depth concentration distributions. The 1 % surface irradiance threshold was derived from Castant et al., 2024 (Castant et al., 2024).

**Table A1.**
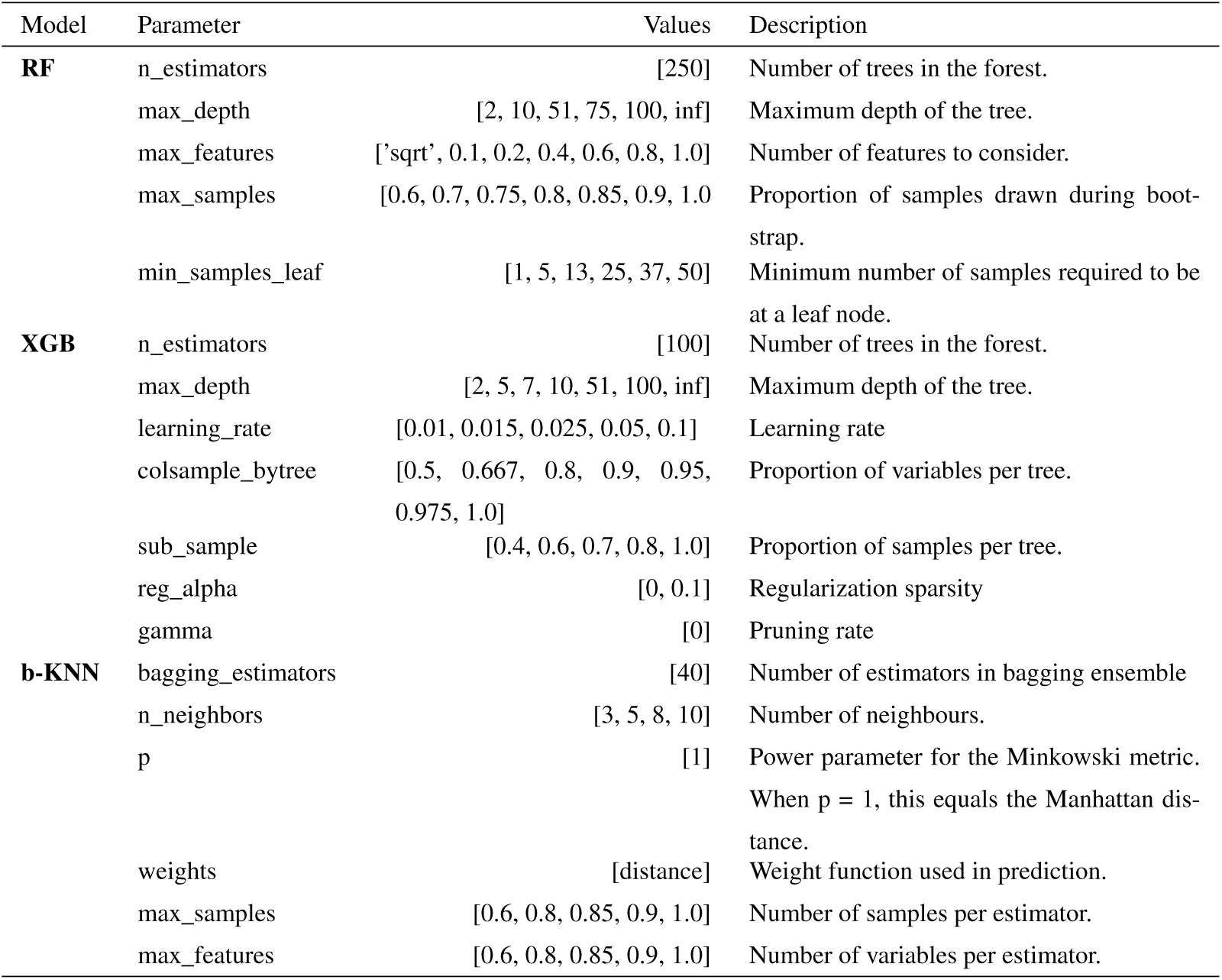
Hyperparameters used in this study. Parameter notation follows SCIKIT-LEARN syntax.

| Model | Parameter | Values | Description |
| --- | --- | --- | --- |
| **RF** | n_estimators | [250] | Number of trees in the forest. |
|  | max_depth | [2, 10, 51, 75, 100, inf] | Maximum depth of the tree. |
|  | max_features | ['sqrt', 0.1, 0.2, 0.4, 0.6, 0.8, 1.0] | Number of features to consider. |
|  | max_samples | [0.6, 0.7, 0.75, 0.8, 0.85, 0.9, 1.0] | Proportion of samples drawn during bootstrap. |
|  | min_samples_leaf | [1, 5, 13, 25, 37, 50] | Minimum number of samples required to be at a leaf node. |
| **XGB** | n_estimators | [100] | Number of trees in the forest. |
|  | max_depth | [2, 5, 7, 10, 51, 100, inf] | Maximum depth of the tree. |
|  | learning_rate | [0.01, 0.015, 0.025, 0.05, 0.1] | Learning rate |
|  | colsample_bytree | [0.5, 0.667, 0.8, 0.9, 0.95, 0.975, 1.0] | Proportion of variables per tree. |
|  | sub_sample | [0.4, 0.6, 0.7, 0.8, 1.0] | Proportion of samples per tree. |
|  | reg_alpha | [0, 0.1] | Regularization sparsity |
|  | gamma | [0] | Pruning rate |
| **b-KNN** | bagging_estimators | [40] | Number of estimators in bagging ensemble |
|  | n_neighbors | [3, 5, 8, 10] | Number of neighbours. |
| | p | [1] | Power parameter for the Minkowski metric.<br>When $p = 1$, this equals the Manhattan distance. |
|  | weights | [distance] | Weight function used in prediction. |
|  | max_samples | [0.6, 0.8, 0.85, 0.9, 1.0] | Number of samples per estimator. |
|  | max_features | [0.6, 0.8, 0.85, 0.9, 1.0] | Number of variables per estimator. |

**Table A2.** Top species percentage contributions to globally integrated abundance. Brackets denote 95th percentile confidence intervals. Species with at least 1 % contribution are listed, for the abundances of all species see Data S2.

| species | contribution (%) |
| --- | --- |
| *Gephyrocapsa huxleyi* | 28.5 [12.3, 46.4] |
| *Gephyrocapsa oceanica* | 7.6 [2.1, 16.0] |
| *Gephyrocapsa ericsonii* | 7.1 [2.4, 14.3] |
| *Florisphaera profunda* | 6.7 [2.3, 13.2] |
| *Umbellosphaera irregularis* | 6.2 [2.7, 11.4] |
| *Oolithotus antillarum* | 3.6 [0.8, 8.5] |
| *Umbellosphaera tenuis* | 2.8 [0.7, 6.4] |
| *Gladiolithus flabellatus* | 2.2 [1.0, 4.1] |
| *Calciopappus caudatus* | 2.1 [0.5, 5.0] |
| *Syracosphaera molischii* | 2.1 [0.7, 4.3] |
| *Ophiaster hydroideus* | 1.9 [0.5, 4.1] |
| *Corisphaera gracilis* HOL | 1.6 [0.6, 3.3] |
| *Gephyrocapsa muelleriae* | 1.6 [0.6, 3.1] |
| *Umbilicosphaera hulburtiana* | 1.5 [0.6, 2.9] |
| *Acanthoica quattropsina* | 1.2 [0.3, 2.9] |
| *Coccolithus pelagicus* | 1.2 [0.2, 3.2] |
| *Acanthoica acanthifera* | 1.2 [0.1, 3.5] |
| *Calciosolenia brasiliensis* | 1.1 [0.3, 2.6] |
| *Palusphaera vandellii* | 1.1 [0.2, 2.7] |
| *Syracosphaera mediterranea* | 1.0 [0.4, 1.9] |
| *Calcidiscus leptoporus* | 1.0 [0.4, 1.9] |
| *Calciosolenia murrayi* | 1.0 [0.1, 2.5] |

**Table A3.** Top species C_IC_ concentrations and contributions to global stocks of the different biomes. “Other" includes all species considered in this study aside from *C. leptoporus*, *C. pelagicus*, *G. huxleyi* and *F. profunda*. Biomes are defined based on Fay and McKinley (Fay and McKinley, 2014).

| Biome | Species | Concentration ($\mu\text{mol/m}^3$) | Contribution (%) |
| --- | --- | --- | --- |
| Global | *Calcidiscus leptoporus* | 1.1 [0.5, 2.0] | 11.7 [4.7, 21.7] |
|  | *Coccolithus pelagicus* | 5.4 [1.4, 11.3] | 24.7 [5.7, 48.0] |
|  | *Gephyrocapsa huxleyi* | 1.0 [0.3, 1.5] | 7.2 [2.4, 14.7] |
|  | *Florisphaera profunda* | 1.3 [0.5, 1.9] | 13.0 [4.7, 24.6] |
|  | other | 3.7 [1.5, 7.4] | 43.3 [28.7, 58.6] |
| Ice | *Calcidiscus leptoporus* | 0.0 [0.0, 0.0] | 0.1 [0.0, 0.3] |
|  | *Coccolithus pelagicus* | 7.1 [2.0, 12.5] | 75.7 [41.2, 93.3] |
|  | *Gephyrocapsa huxleyi* | 0.1 [0.0, 0.2] | 3.6 [0.7, 10.6] |
|  | *Florisphaera profunda* | 0.0 [0.0, 0.0] | 0.0 [0.0, 0.0] |
|  | other | 0.4 [0.0, 2.7] | 20.6 [4.8, 53.6] |
| Subpolar | *Calcidiscus leptoporus* | 1.6 [0.7, 3.3] | 12.9 [3.3, 31.0] |
|  | *Coccolithus pelagicus* | 15.5 [4.8, 33.9] | 64.3 [29.5, 86.3] |
|  | *Gephyrocapsa huxleyi* | 3.0 [1.1, 4.5] | 16.8 [4.8, 37.7] |
|  | *Florisphaera profunda* | 0.0 [0.0, 0.0] | 0.0 [0.0, 0.2] |
|  | other | 0.7 [0.2, 2.4] | 6.0 [2.2, 13.1] |
| Subtropical | *Calcidiscus leptoporus* | 2.6 [1.3, 4.8] | 20.2 [7.5, 36.7] |
| Seasonally Stratified | *Coccolithus pelagicus* | 4.8 [0.6, 13.7] | 30.8 [3.7, 63.3] |
|  | *Gephyrocapsa huxleyi* | 1.5 [0.5, 2.2] | 8.7 [2.6, 18.7] |
|  | *Florisphaera profunda* | 1.0 [0.4, 1.4] | 6.4 [2.3, 12.9] |
|  | other | 4.5 [1.7, 8.6] | 33.9 [17.7, 51.7] |
| Subtropical | *Calcidiscus leptoporus* | 1.0 [0.5, 1.8] | 10.7 [4.6, 18.8] |
| Permanently Stratified | *Coccolithus pelagicus* | 0.4 [0.1, 0.9] | 3.5 [0.5, 9.3] |
|  | *Gephyrocapsa huxleyi* | 0.5 [0.1, 0.8] | 3.9 [1.0, 8.5] |
|  | *Florisphaera profunda* | 2.4 [0.8, 3.5] | 20.5 [8.6, 34.9] |
|  | other | 5.7 [2.4, 10.7] | 61.4 [48.5, 73.9] |
| Equatorial | *Calcidiscus leptoporus* | 0.4 [0.2, 0.8] | 8.2 [3.2, 15.6] |
|  | *Coccolithus pelagicus* | 0.0 [0.0, 0.0] | 0.1 [0.0, 0.3] |
|  | *Gephyrocapsa huxleyi* | 0.1 [0.0, 0.2] | 1.7 [0.4, 3.9] |
|  | *Florisphaera profunda* | 1.7 [0.5, 2.3] | 21.5 [7.9, 38.7] |
|  | other | 3.4 [1.3, 7.5] | 68.5 [52.3, 82.8] |

**Table A4.**
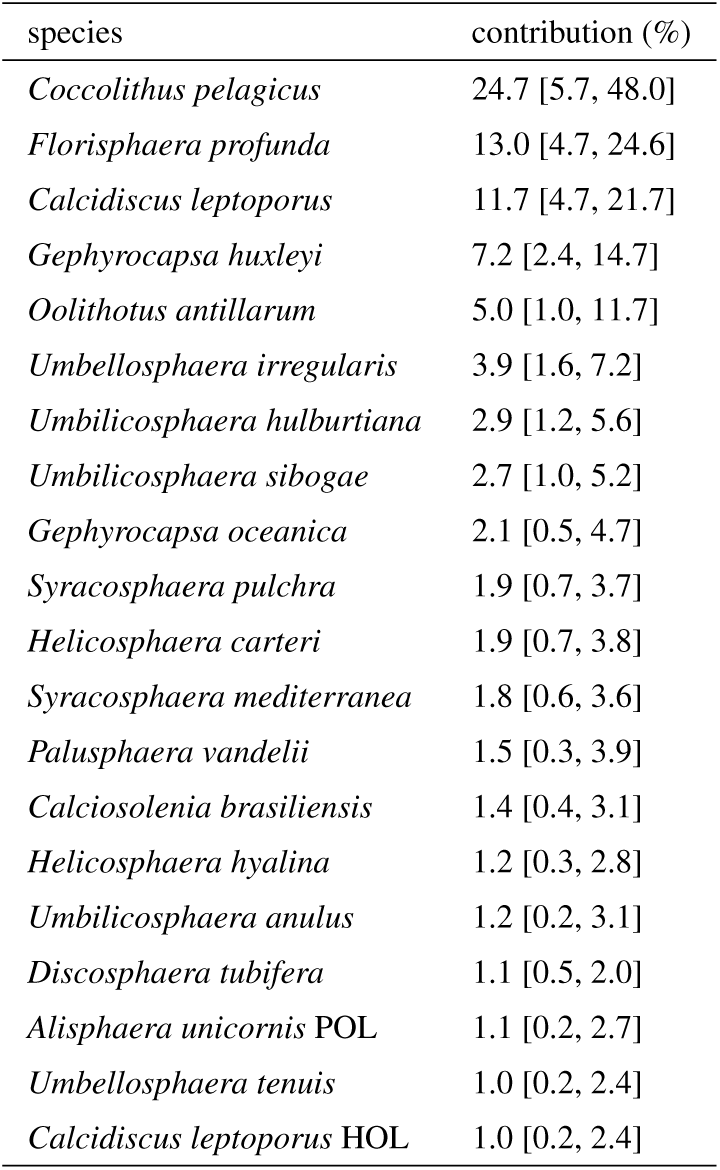
Top species C_IC_ contributions to globally integrated C_IC_. Species with at least 1 % contribution are listed, for the C_IC_ contributions of all species see Data S3.

| species | contribution (%) |
| --- | --- |
| *Coccolithus pelagicus* | 24.7 [5.7, 48.0] |
| *Florisphaera profunda* | 13.0 [4.7, 24.6] |
| *Calcidiscus leptoporus* | 11.7 [4.7, 21.7] |
| *Gephyrocapsa huxleyi* | 7.2 [2.4, 14.7] |
| *Oolithotus antillarum* | 5.0 [1.0, 11.7] |
| *Umbellosphaera irregularis* | 3.9 [1.6, 7.2] |
| *Umbilicosphaera hulburtiana* | 2.9 [1.2, 5.6] |
| *Umbilicosphaera sibogae* | 2.7 [1.0, 5.2] |
| *Gephyrocapsa oceanica* | 2.1 [0.5, 4.7] |
| *Syracosphaera pulchra* | 1.9 [0.7, 3.7] |
| *Helicosphaera carteri* | 1.9 [0.7, 3.8] |
| *Syracosphaera mediterranea* | 1.8 [0.6, 3.6] |
| *Palusphaera vandellii* | 1.5 [0.3, 3.9] |
| *Calciosolenia brasiliensis* | 1.4 [0.4, 3.1] |
| *Helicosphaera hyalina* | 1.2 [0.3, 2.8] |
| *Umbilicosphaera anulus* | 1.2 [0.2, 3.1] |
| *Discosphaera tubifera* | 1.1 [0.5, 2.0] |
| *Alisphaera unicornis* POL | 1.1 [0.2, 2.7] |
| *Umbellosphaera tenuis* | 1.0 [0.2, 2.4] |
| *Calcidiscus leptoporus* HOL | 1.0 [0.2, 2.4] |

### Author contributions

Conceptualization: JdV, FMM; Data curation: JdV, NAW; Formal analysis: JdV; Funding Acquisition: FMM, JdV, LJW, AJP; Methodology: JdV, LJW; Software: JdV, NAW, LJW; JdV, NAW; Writing – original draft: JdV, FMM, LJW, AJP, NAW.

### Competing interests

There are no competing interests to declare.

## Acknowledgements

This work was supported by funding from the UK Research and Innovation Natural Environment Research Council grant CoccoTrait, NE/X001261/1 (FMM, JdV, AJP, LJW); UK Research and Innovation Natural Environment Research Council GW4+ DTP studentship, NE/L002434/1 (JdV); European Union under grant agreement no. 101083922, OceanICU (AJP); and the UK Research and Innovation (UKRI) under the UK government’s Horizon Europe funding guarantee, grant number 10054454 (AJP). Views and opinions expressed are however those of the author(s) only and do not necessarily reflect those of the European Union or European Research Executive Agency. Neither the European Union nor the granting authority can be held responsible for them.

This work was carried out using the computational facilities of the Advanced Computing Research Centre, University of Bristol - http://www.bristol.ac.uk/acrc/. We thank all researchers and technical staff involved in field observations and acknowledge their efforts in making this data publicly available. We also thank the developers who maintain the open-source software used to perform this research (e.g. Unix, Python, scikit-learn).

## References

Agusti, S., González-Gordillo, J. I., Vaqué, D., Estrada, M., Cerezo, M. I., Salazar, G., Gasol, J. M., and Duarte, C. M.: Ubiquitous healthy diatoms in the deep sea confirm deep carbon injection by the biological pump, Nature Communications, 6, 7608, 10.1038/ncomms8608, 2015.

Balch, W. M.: The ecology, biogeochemistry, and optical properties of coccolithophores, Annual review of marine science, 10, 71–98, 2018.

Balch, W. M. and Mitchell, C.: Remote sensing algorithms for particulate inorganic carbon (PIC) and the global cycle of PIC, Earth-Science Reviews, 239, 104 363, 2023.

Balch, W. M., Gordon, H. R., Bowler, B., Drapeau, D., and Booth, E.: Calcium carbonate measurements in the surface global ocean based on Moderate-Resolution Imaging Spectroradiometer data, Journal of Geophysical Research: Oceans, 110, 2005.

Benedetti, F., Gruber, N., and Vogt, M.: Global gradients in species richness of marine plankton functional groups, Journal of Plankton Research, 45, 832–852, 10.1093/plankt/fbad056, 2023.

Boudreau, B. P., Middelburg, J. J., and Luo, Y.: The role of calcification in carbonate compensation, Nature Geoscience, 11, 894–900, 2018.

Breiman, L.: Random Forests, Machine Learning, 45, 5–32, 10.1023/A:1010933404324, 2001.

Broullón, D., Pérez, F. F., Velo, A., Hoppema, M., Olsen, A., Takahashi, T., Key, R. M., Tanhua, T., González-Dávila, M., Jeansson, E., Kozyr, A., Van Heuven, A. C., et al.: A global monthly climatology of total alkalinity: A neural network approach, Earth System Science Data, 11, 1109–1127, 10.5194/essd-11-1109-2019, 2019.

Broullón, D., Pérez, F. F., Velo, A., Hoppema, M., Olsen, A., Takahashi, T., Key, R. M., Tanhua, T., Santana-Casiano, J. M., and Kozyr, A.: A global monthly climatology of oceanic total dissolved inorganic carbon: A neural network approach, Earth System Science Data, 12, 1725–1743, 10.5194/essd-12-1725-2020, 2020.

Buitenhuis, E. T., Vogt, M., Moriarty, R., Bednaršek, N., Doney, S. C., Leblanc, K., Le Quéré, C., Luo, Y.-W., O’Brien, C., O’Brien, T., et al.: MAREDAT: towards a world atlas of MARine Ecosystem DATa, Earth System Science Data, 5, 227–239, 2013.

Castant, J., Vantrepotte, V., Frouin, R., and Beaugrand, G.: Comprehensive gridded dataset of photosynthetically active radiation in the upper ocean from 1958 to 2022, Remote Sensing of Environment, 311, 14 305, 10.1016/j.rse.2024.14305, 2024.

Chen, T. and Guestrin, C.: XGBoost: A scalable tree boosting system, in: Proceedings of the ACM SIGKDD International Conference on Knowledge Discovery and Data Mining, pp. 785–794, 2016.

Cover, T. and Hart, P.: Nearest neighbor pattern classification, IEEE Transactions on Information Theory, 13, 21–27, 1967.

Daniels, C. J., Sheward, R. M., and Poulton, A. J.: Biogeochemical implications of comparative growth rates of Emiliania huxleyi and Coccolithus species, Biogeosciences, 11, 6915–6925, 2014.

Daniels, C. J., Poulton, A. J., Young, J. R., Esposito, M., Humphreys, M. P., Ribas-Ribas, M., Tynan, E., and Tyrrell, T.: Species-specific calcite production reveals Coccolithus pelagicus as the key calcifier in the Arctic Ocean, Marine Ecology Progress Series, 555, 29–47, 2016.

de Vries, J., Monteiro, F., Wheeler, G., Poulton, A., Godrijan, J., Cerino, F., Malinverno, E., Langer, G., and Brownlee, C.: Haplo-diplontic life cycle expands coccolithophore niche, Biogeosciences, 18, 1161–1184, 2021.

de Vries, J., Poulton, A. J., Young, J. R., Monteiro, F. M., Sheward, R. M., Johnson, R., Hagino, K., Ziveri, P., and Wolf, L. J.: CASCADE: Dataset of extant coccolithophore size, carbon content and global distribution, Scientific data, 11, 920, 2024.

de Vries, J., Wiseman, N., and Wolf, L. J.: nanophyto/Abil: v25.08.06, 10.5281/zenodo.16886568, 2025a.

de Vries, J., Wiseman, N. A., and Wolf, L. J.: Abil: A Python package for the interpolation of aquatic biogeochemical datasets, Journal of Open Source Software, 10, 8755, 2025b.

Fay, A. R. and McKinley, G. A.: Global open-ocean biomes: Mean and temporal variability, Earth System Science Data, 6, 273–284, 10.5194/essd-6-273-2014, 2014.

Flynn, K. J., Stoecker, D. K., Mitra, A., Raven, J. A., Glibert, P. M., Hansen, P. J., Granéli, E., and Burkholder, J. M.: Misuse of the phytoplankton-zooplankton dichotomy: The need to assign organisms as mixotrophs within plankton functional types, Journal of Plankton Research, 35, 3–11, 10.1093/plankt/fbs072, 2013.

Gafar, N., Eyre, B., and Schulz, K.: Particulate inorganic to organic carbon production as a predictor for coccolithophorid sensitivity to ongoing ocean acidification, Limnology and Oceanography Letters, 4, 62–70, 2019.

Gibbs, S. J., Bown, P. R., Ward, B. A., Alvarez, S. A., Kim, H., Archontikis, O. A., Sauterey, B., Poulton, A. J., Wilson, J., and Ridgwell, A.: Algal plankton turn to hunting to survive and recover from end-Cretaceous impact darkness, Science Advances, 6, eabc9123, 10.1126/sciadv.abc9123, 2020.

Guerreiro, C. V., Baumann, K.-H., Brummer, G.-J. A., Fischer, G., Korte, L. F., Merkel, U., Sá, C., De Stigter, H., and Stuut, J.-B. W.: Coccolithophore fluxes in the open tropical North Atlantic: influence of thermocline depth, Amazon water, and Saharan dust, Biogeosciences, 14, 4577–4599, 2017.

Géron, A.: Hands-on Machine Learning with Scikit-Learn, Keras, and TensorFlow, O’Reilly Media, Inc., 2022.

Han, Y., Steiner, Z., Cao, Z., Fan, D., Chen, J., Yu, J., and Dai, M.: Coccolithophore abundance and production and their impacts on particulate inorganic carbon cycling in the western North Pacific, Biogeosciences, 22, 3681–3697, 2025.

Hopkins, J., Henson, S. A., Painter, S. C., Tyrrell, T., and Poulton, A. J.: Phenological characteristics of global coccolithophore blooms, Global Biogeochemical Cycles, 29, 239–253, 2015.

Hopkins, J., Henson, S. A., Poulton, A. J., and Balch, W. M.: Regional characteristics of the temporal variability in the global particulate inorganic carbon inventory, Global Biogeochemical Cycles, 33, 1328–1338, 2019.

Knecht, N. S., Benedetti, F., Hofmann Elizondo, U., Bednaršek, N., Chaabane, S., de Weerd, C., Peijnenburg, K. T., Schiebel, R., and Vogt, M.: The impact of zooplankton calcifiers on the marine carbon cycle, Global Biogeochemical Cycles, 37, e2022GB007 685, 2023.

Kostadinov, T. S., Milutinović, S., Marinov, I., and Cabré, A.: Carbon-based phytoplankton size classes retrieved via ocean color estimates of the particle size distribution, Ocean Science, 12, 561–575, 2016.

Kottmeier, D. M., Chrachri, A., Langer, G., Helliwell, K. E., Wheeler, G. L., and Brownlee, C.: Reduced H+ channel activity disrupts pH homeostasis and calcification in coccolithophores at low ocean pH, Proceedings of the National Academy of Sciences, 119, e2118009 119, 2022.

Kruijt, A. L., van Dijk, R., Sulpis, O., Beaufort, L., Lassus, G., Brummer, G.-J., van der Burg, A. D., Cala, B. A., Ourradi, Y., Peijnenburg, K. T., et al.: The contributions of various calcifying plankton to the South Atlantic calcium carbonate stock, Biogeosciences, 23, 531–563, 2026.

Krumhardt, K. M., Lovenduski, N. S., Iglesias-Rodriguez, M. D., and Kleypas, J. A.: Coccolithophore growth and calcification in a changing ocean, Progress in oceanography, 159, 276–295, 2017.

Krumhardt, K. M., Lovenduski, N. S., Long, M. C., Lévy, M., Lindsay, K., Moore, J. K., and Nissen, C.: Coccolithophore growth and calcification in an acidified ocean: Insights from community Earth system model simulations, Journal of Advances in Modeling Earth Systems, 11, 1418–1437, 2019.

Krumhardt, K. M., Long, M. C., Lindsay, K., and Levy, M. N.: Southern Ocean calcification controls the global distribution of alkalinity, Global Biogeochemical Cycles, 34, e2020GB006 727, 2020.

Kwon, E. Y., Dunne, J. P., and Lee, K.: Biological export production controls upper ocean calcium carbonate dissolution and CO2 buffer capacity, Science Advances, 10, eadl0779, 2024.

Le Quéré, C., Buitenhuis, E. T., Moriarty, R., Alvain, S., Aumont, O., Bopp, L., Chollet, S., Enright, C., Franklin, D. J., Geider, R. J., et al.: Role of zooplankton dynamics for Southern Ocean phytoplankton biomass and global biogeochemical cycles, Biogeosciences, 13, 4111–4133, 2016.

Litchman, E. and Klausmeier, C. A.: Trait-based community ecology of phytoplankton, Annual review of ecology, evolution, and systematics, 39, 615–639, 2008.

Meyer, H. and Pebesma, E.: Predicting into unknown space? Estimating the area of applicability of spatial prediction models, Methods in Ecology and Evolution, 12, 1620–1633, 2021.

Monteiro, F. M., Bach, L. T., Brownlee, C., Bown, P., Rickaby, R. E., Poulton, A. J., Tyrrell, T., Beaufort, L., Dutkiewicz, S., Gibbs, S., et al.: Why marine phytoplankton calcify, Science Advances, 2, e1501 822, 2016.

Neukermans, G., Bach, L., Butterley, A., Sun, Q., Claustre, H., and Fournier, G.: Quantitative and mechanistic understanding of the open ocean carbonate pump-perspectives for remote sensing and autonomous in situ observation, Earth-Science Reviews, 239, 104 359, 2023.

O’Brien, C. J., Peloquin, J. A., Vogt, M., Heinle, M., Gruber, N., Ajani, P., Andruleit, H., Arístegui, J., Beaufort, L., Estrada, M., et al.: Global marine plankton functional type biomass distributions: coccolithophores, Earth System Science Data, 5, 259–276, 2013.

O’Brien, C. J., Vogt, M., and Gruber, N.: Global coccolithophore diversity: Drivers and future change, Progress in Oceanography, 140, 27–42, 2016.

Planchat, A., Kwiatkowski, L., Bopp, L., Torres, O., Christian, J. R., Butenschön, M., Lovato, T., Séférian, R., Chamberlain, M. A., Aumont, O., Watanabe, M., Yamamoto, A., Yool, A., Ilyina, T., Tsujino, H., Krumhardt, K. M., Schwinger, J., Tjiputra, J., Dunne, J. P., and Stock, C.: The representation of alkalinity and the carbonate pump from CMIP5 to CMIP6 Earth system models and implications for the carbon cycle, Biogeosciences, 20, 1195–1257, 10.5194/bg-20-1195-2023, 2023.

Planchat, A., Bopp, L., Kwiatkowski, L., and Torres, O.: The carbonate pump feedback on alkalinity and the carbon cycle in the 21st century and beyond, Earth System Dynamics, 15, 565–588, 2024.

Poulton, A. J., Adey, T. R., Balch, W. M., and Holligan, P. M.: Relating coccolithophore calcification rates to phytoplankton community dynamics: Regional differences and implications for carbon export, Deep Sea Research Part II: Topical Studies in Oceanography, 54, 538–557, 2007.

Poulton, A. J., Holligan, P. M., Charalampopoulou, A., and Adey, T. R.: Coccolithophore ecology in the tropical and subtropical Atlantic Ocean: New perspectives from the Atlantic meridional transect (AMT) programme, Progress in Oceanography, 158, 150–170, 2017.

Reagan, J. R., Boyer, T. P., García, H. E., Locarnini, R. A., Baranova, O. K., Bouchard, C., Cross, S. L., Mishonov, A. V., Paver, C. R., Seidov, D., Wang, Z., and Dukhovskoy, D.: World Ocean Atlas 2023, World Ocean Atlas, 2024.

Rideout, J. R., Caporaso, G., Bolyen, E., McDonald, D., Baeza, Y. V., Alastuey, J. C., Pitman, A., Morton, J., Navas, J., Gorlick, K., Debelius, J., Xu, Z., llcooljohn, adamrp, Shorenstein, J., Luce, L., Treuren, W. V., charudatta navare, Gonzalez, A., Brislawn, C. J., Patena, W., Schwarzberg, K., teravest, Reeder, J., shiffer1, Sfiligoi, I., nbresnick, Zhu, Q., Murray, D. K. D., and Sharma, K.: biocore/scikit-bio: scikit-bio 0.5.9: Maintenance release, 10.5281/zenodo.8209901, 2023.

Ridgwell, A. and Zeebe, R. E.: The role of the global carbonate cycle in the regulation and evolution of the Earth system, Earth and Planetary Science Letters, 234, 299–315, 2005.

Rigual Hernández, A. S., Trull, T. W., Nodder, S. D., Flores, J. A., Bostock, H., Abrantes, F., Eriksen, R. S., Sierro, F. J., Davies, D. M., Ballegeer, A.-M., et al.: Coccolithophore biodiversity controls carbonate export in the Southern Ocean, Biogeosciences, 17, 245–263, 2020.

Seifert, M., Nissen, C., Rost, B., and Hauck, J.: Cascading effects augment the direct impact of CO2 on phytoplankton growth in a biogeochemical model, Elem Sci Anth, 10, 00 104, 2022.

Sheward, R. M., Poulton, A. J., Young, J. R., de Vries, J., Monteiro, F. M., and Herrle, J. O.: Cellular morphological trait dataset for extant coccolithophores from the atlantic ocean, Scientific Data, 11, 720, 2024.

Tittensor, D. P., Mora, C., Jetz, W., Lotze, H. K., Ricard, D., Vanden Berghe, E., and Worm, B.: Global patterns and predictors of marine biodiversity across taxa, Nature, 466, 1098–1101, 10.1038/nature09329, 2010.

Varoquaux, G., Buitinck, L., Louppe, G., Grisel, O., Pedregosa, F., and Mueller, A.: Scikit-learn: Machine learning without learning the machinery, GetMobile: Mobile Computing and Communications, 19, 29–33, 2015.

Villiot, N., Poulton, A. J., Butcher, E. T., Daniels, L. R., and Coggins, A.: Allometry of carbon and nitrogen content and growth rate in a diverse range of coccolithophores, Journal of Plankton Research, 43, 511–526, 2021.

Young, J. R.: Functions of coccoliths, in: Coccolithophores, edited by Winter, A. and Siesser, W. G., pp. 63–82, Cambridge University Press, Cambridge, 1994.

Young, J. R., Bown, P. R., and Lees, J. A.: Nannotax3 website, Online; International Nannoplankton Association, https://www.mikrotax.org/Nannotax3, accessed: 2025-08-29, 2022.

Zhang, Z., Zhang, S., Behrenfeld, M. J., Chen, P., Jamet, C., Di Girolamo, P., Dionisi, D., Hu, Y., Lu, X., Pan, Y., et al.: Combining deep learning with physical parameters in POC and PIC inversion from spaceborne lidar CALIOP, ISPRS Journal of Photogrammetry and Remote Sensing, 212, 193–211, 2024.

Ziveri, P., de Bernardi, B., Baumann, K.-H., Stoll, H. M., and Mortyn, P. G.: Sinking of coccolith carbonate and potential contribution to organic carbon ballasting in the deep ocean, Deep Sea Research Part II: Topical Studies in Oceanography, 54, 659–675, 2007.

Ziveri, P., Gray, W. R., Anglada-Ortiz, G., Manno, C., Grelaud, M., Incarbona, A., Rae, J. W. B., Subhas, A. V., Pallacks, S., White, A., et al.: Pelagic calcium carbonate production and shallow dissolution in the North Pacific Ocean, Nature Communications, 14, 805, 2023.

Ziveri, P., Langer, G., Chaabane, S., de Vries, J., Gray, W. R., Keul, N., Hatton, I. A., Manno, C., Norris, R., Pallacks, S., et al.: Calcifying plankton: From biomineralization to global change, Science, 390, eadq8520, 2025.

